# High-resolution transcriptomics of bovine purified protein derivative-stimulated peripheral blood from cattle infected with *Mycobacterium bovis* across an experimental time course

**DOI:** 10.1101/2022.04.19.488769

**Authors:** Carolina N. Correia, Gillian P. McHugo, John A. Browne, Kirsten E. McLoughlin, Nicolas C. Nalpas, David A. Magee, Adam O. Whelan, Bernardo Villarreal-Ramos, H. Martin Vordermeier, Eamonn Gormley, Stephen V. Gordon, David E. MacHugh

## Abstract

**Objectives:** Improved bovine tuberculosis (bTB) diagnostics with higher sensitivity and specificity are urgently required. A better understanding of the peripheral blood transcriptional response of *Mycobacterium bovis*-infected animals after bovine purified protein derivative (PPD-b) stimulation of whole blood—an important component of current bTB diagnostics—will provide new information for development of better diagnostics.

**Methods:** RNA sequencing (RNA-seq) was used to study the peripheral blood transcriptome after stimulation with PPD-b across four time points (-1 wk pre-infection, and +1 wk, +2 wk, and +10 wk post-infection) from a 14-week *M. bovis* infection time course experiment with ten age-matched Holstein-Friesian cattle.

**Results:** In vitro PPD-b stimulation of peripheral blood from *M. bovis*-infected and non-infected cattle elicited a strong transcriptional response. Comparison of PPD-b stimulated, and unstimulated samples revealed higher expression of genes encoding cytokine receptors, transcription factors, and interferon-inducible proteins. Lower expression was seen for genes encoding proteins involved in antimicrobial activity, C-type lectin receptors, inhibition of signal transduction, and genes encoding metal ion transporters.

**Conclusions:** A transcriptional signature associated with the peripheral blood response to PPD-b stimulation consisting of 170 genes was identified exclusively in the post-infection time points. Therefore, this represents a panel of potential biomarkers of *M. bovis* infection.

## 1. Introduction

The impacts of *Mycobacterium bovis* infection, the primary cause of bovine tuberculosis (bTB), involve the zoonotic risk of human infection, losses in animal productivity, and the economic consequences resulting from herd restrictions, disruptions to trade and reduced agricultural productivity [1–5]. In Ireland, approximately 30,000 bTB-infected cattle have been culled on average every year since the mid-1960s [6, 7] with recent figures indicating annual costs of up to €97 million [8]. In addition, on a global scale, a longstanding and conservative estimate suggests that bTB costs at least $3 billion annually and imposes a large financial burden on farmers with infected herds, particularly in developing countries [9, 10].

The current strategy underpinning bTB control is the “test and slaughter” programme. The cornerstone of this strategy in Ireland and the UK is the single intradermal comparative tuberculin test (SICTT), whereby infected animals are identified based on their comparative immunological response to intradermal injections with crude preparations of mycobacterial antigens (purified protein derivative or PPD) [11, 12]. PPD from two different species of mycobacteria are used in the SICTT: PPD-bovine (PPD-b) derived from heat-killed cultures of *M. bovis* strain AN5, and a comparative PPD from *M. avium*, a common environmental mycobacterium (PPD-avian, or PPD-a), which is used to increase the specificity of the test. Due to a lack of absolute animal test sensitivity (typically 70-90%, depending on the stage of the disease, the type of test used, and the severity of test interpretation) the test performance fails to detect all infected animals [13, 14]. In Ireland and the UK, an ancillary whole-blood diagnostic test is routinely used in parallel—the interferon-γ (IFN-γ) release assay (IGRA)—that has estimated sensitivity and specificity of approximately 85% and 95%, respectively [14, 15]. It is widely recognised that improved bTB diagnostics with higher sensitivity and specificity are urgently required to augment existing tests and refine and enhance current bTB control and eradication programmes around the world [4, 12, 16].

Peripheral blood gene expression studies can offer a detailed snapshot of genome-wide changes in host mRNA transcript abundance and have, therefore, been widely used for development of diagnostic and prognostic biomarkers of the host immune response under pathological conditions, including for human tuberculosis caused by infection with *Mycobacterium tuberculosis* [17–21]. In this regard, functional genomics technologies, such as gene expression microarrays and RNA sequencing (RNA-seq) have also been used to identify and characterise peripheral blood RNA biomarkers and biosignatures of *M. bovis* infection and bTB disease [22–31].

Understanding more about the transcriptional response of non-infected and *M. bovis*-infected animals after PPD-b stimulation—an important component of current bTB diagnostic tests—will provide a better understanding of the host immune response and new information for development of better diagnostics [10, 16, 32]. In the present study, RNA-seq was used to study the bovine peripheral blood transcriptome after stimulation with PPD-b across four time points from a 14-week *M. bovis* infection time course experiment with ten animals. We evaluated host peripheral blood transcriptional responses to PPB-b stimulation across one time point pre-infection (-1 wk) and three time-points (+1 wk, +2 wk, and + 10 wk) post-infection with *M. bovis* and identified pathways enriched after PPB-b stimulation in non-infected and *M. bovis*-infected cattle.

## 2. Materials and methods

### 2.1. Ethics statement

Animal experimental work was carried out according to the UK Animal (Scientific Procedures) Act 1986. The study protocol was approved by the Animal Health and Veterinary Laboratories Agency (AHVLA-Weybridge, UK), now the Animal & Plant Health Agency (APHA), Animal Use Ethics Committee (UK Home Office PCD Number 70/6905).

### 2.2. Animal infection time course experiment

Ten age-matched male Holstein-Friesian cattle (4-7 months old) were sourced from farms known to be free of bTB. The animals used for the experimental work detailed in this study were the naïve control group for a larger vaccine efficacy study that has been described previously [33, 34]. Briefly, four treatment groups consisting of ten animals each were immunised with BCG and boosted with adenovirus or protein-based vaccines at eight weeks post-BCG vaccination. An additional ten-animal group, the naïve controls, did not receive any immunization. Following this, at 12 weeks post-BCG vaccination, the vaccinated and non-vaccinated control animals were infected via the endobronchial route with *M. bovis* AF2122/97 [35] at 2 × 10^3^ CFU per animal, as previously described by Whelan, et al. [36]; the animals were then monitored across a 14-week time course. The group of ten naïve animals were subdivided into two groups of five due to housing space requirements. The first group of five naïve animals (animal IDs starting with 65 – see Supplementary Table 1) underwent complete sampling between May and August 2011, whereas the second naïve subgroup (animal IDs starting with 66 – see Supplementary Table 1) were sampled between August and November 2011. **Fig. 1** details the experimental schedule used by Dean, et al. [33] and shows the sampling time points from the ten non-vaccinated control cattle used for the research work described here.

**Fig. 1:**
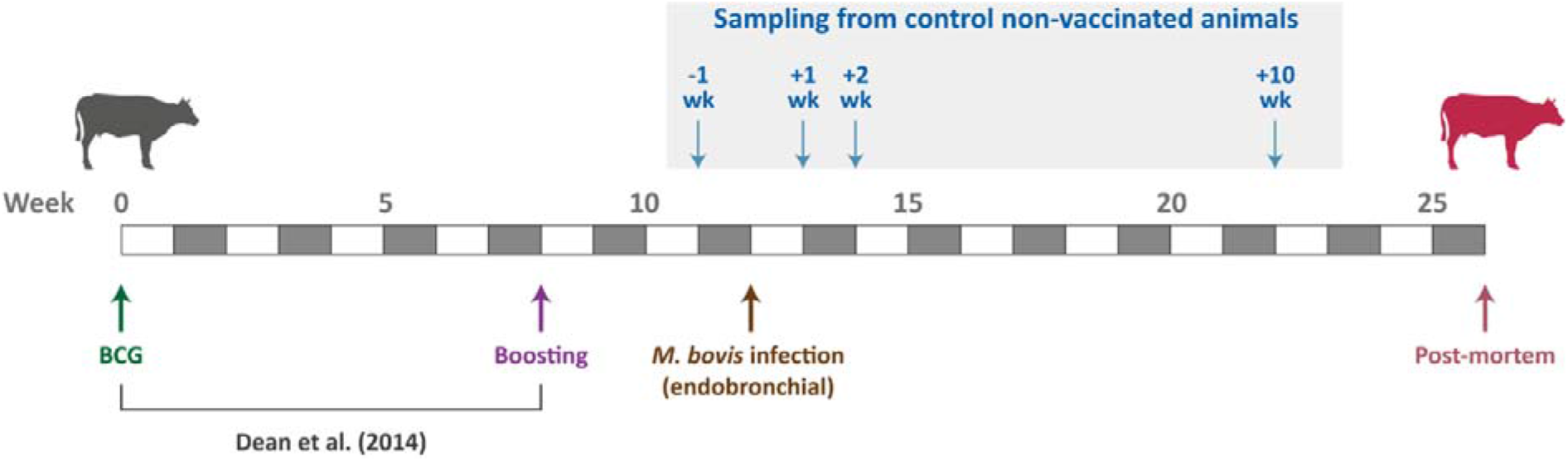
Schedule for the *M. bovis* infection time course experiment. Sampling time points for the ten nonLvaccinated control cattle used are shown. Figure modified from McLoughlin, *et al*. [29].

### 2.3. Blood collection and PPD-b stimulation

Approximately 3 ml of whole blood were collected into heparin-coated Vacutainer^®^ blood collection tubes (BD Diagnostics, Oxford, UK) from all ten non-vaccinated control animals at -1 wk pre-infection; and +1 wk, +2 wk, and +10 wk post-infection (**Fig. 1**), totalling 80 samples. These were used for a) an overnight stimulation with PPD-b at 37°C and b) a control overnight incubation at 37°C without PPD-b stimulation (**Fig. 2A**). Following this, samples were transferred to Tempus^™^ blood RNA tubes (Applied Biosystems^™^/Thermo Fisher Scientific, Warrington, UK). These collection tubes contained 6 ml of a proprietary stabilising reagent that lyses red blood cells, inactivates cellular RNases, and selectively precipitates RNA while genomic DNA and proteins remain in solution. Immediately after the transfer, Tempus^™^ tube samples were vortexed for approximately 10 s to ensure complete red blood cell lysis; lysate samples were then stored at - 80°C.

**Fig. 2:**
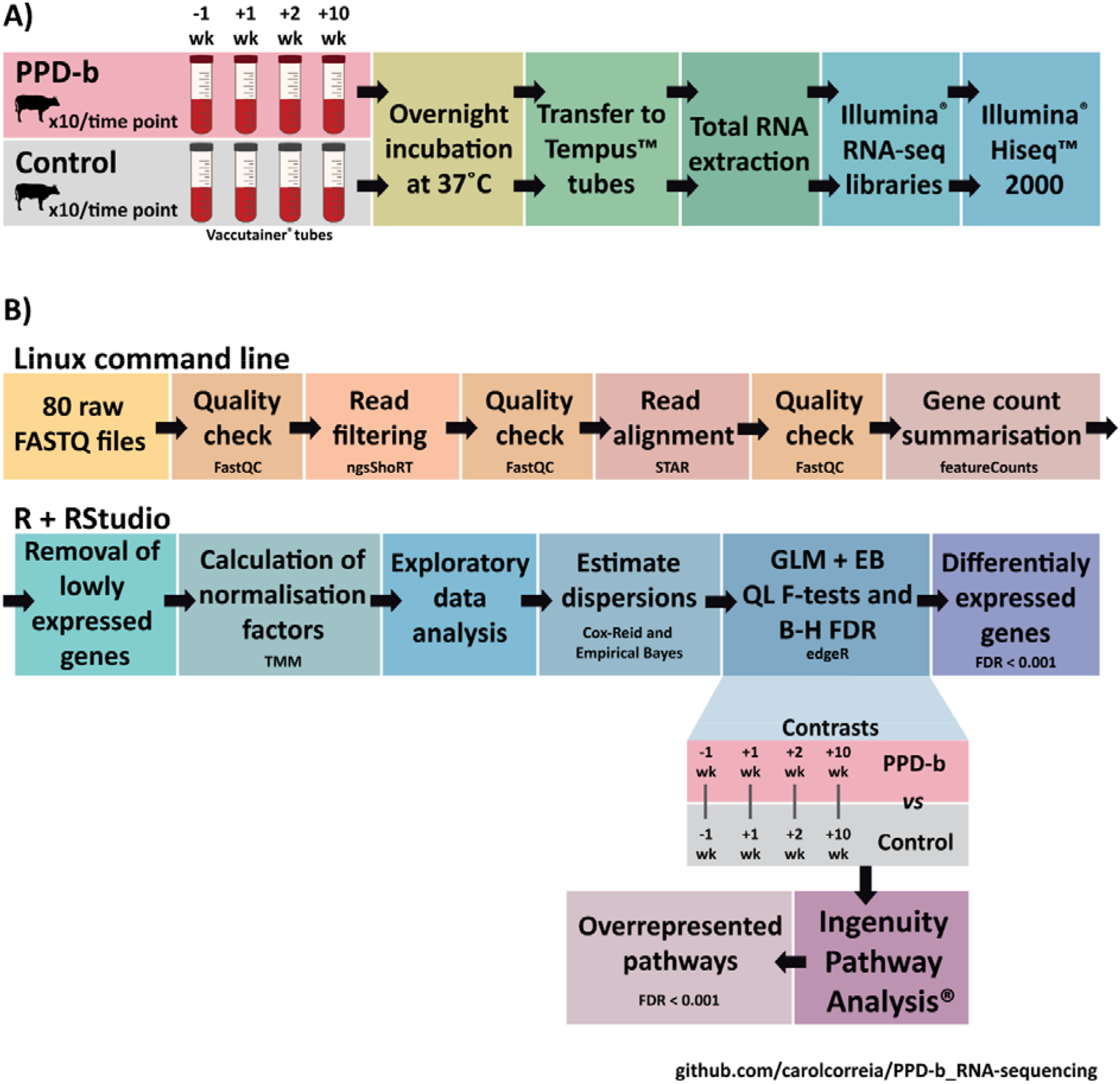
Overview of experimental work and bioinformatics workflow. **A.** PPD-b stimulation experimental design, and RNA-seq library preparation and sequencing. **B.** Data pre-processing and statistical procedures for differential gene expression and overrepresented pathway analyses.

### 2.4. Total RNA extraction and purification

The Tempus^™^ Spin RNA Isolation Kit (Applied Biosystems^™^/Thermo Fisher Scientific) was used for total RNA extraction and purification using the manufacturer’s protocol. Tempus^™^ tube blood lysate samples were thawed at room temperature prior to RNA extraction and purification. Once thawed, for each sample, approximately 3 ml of blood lysate was transferred to a 50 ml plastic centrifuge tube and PBS was added to a final volume of 12 ml. Each sample was then mixed by vortexing for 30 s and then centrifuged at 3,000 × g for 30 min at 4°C. The supernatant was then removed, and the remaining RNA-containing pellet was re-suspended with a brief vortex in 400 μl of the proprietary Tempus^™^ RNA Purification Resuspension Solution. Following this, the re-suspended RNA sample was pipetted into an RNA purification filter inserted into a 1.5 ml microcentrifuge tube for waste collection. The RNA purification filter/microcentrifuge tube was then centrifuged at 16,000 × g for 30 s and the liquid waste and microcentrifuge tube discarded. The RNA purification filter was then placed in a clean microcentrifuge tube, 500 μl of proprietary RNA Purification Wash Solution 1 was added, followed by another centrifugation step at 16,000 × g for 30 s and disposal of the liquid waste and microcentrifuge tube. This step was then repeated using 500 μl of proprietary RNA Purification Wash Solution 2 with a centrifugation step at 16,000 × g for 30 s. A final wash step was then performed with 500 μl of RNA Purification Wash Solution 2 and centrifugation at 16,000 × g for 30 s followed by disposal of the liquid waste and microcentrifuge tube. The RNA purification filter was then placed in a clean microcentrifuge tube and centrifuged at 16,000 × g for 30 s to dry the membrane. The RNA purification filter was then inserted into a clean RNase-free collection microcentrifuge tube and 100 μl of Nucleic Acid Purification Elution Solution was added and incubated for 2 min followed by centrifugation at 16,000 × g for 30 s; the RNA eluate was then pipetted back onto the filter membrane and the centrifugation step was repeated. Approximately 90 μl of the final RNA eluate was then pipetted (avoiding particulate material) into a new RNase-free collection microcentrifuge tube for long-term storage at −80°C.

### 2.5. RNA quality checking and quantification

RNA quantity and quality were checked using a NanoDrop^™^ 1000 spectrophotometer (Thermo Fisher Scientific, Waltham, MA, USA) and an Agilent 2100 Bioanalyzer using an RNA 6000 Nano LabChip kit (Agilent Technologies Ltd., Cork, Ireland), according to the manufacturers’ instructions. Supplementary Table 1 details the RNA integrity number (RIN) and concentration data for all samples.

### 2.6. Strand-specific RNA-seq library preparation and sequencing

One μg of total RNA from each sample was used to prepare individually barcoded strand-specific RNA-seq libraries (Supplementary Table 1). Two rounds of poly(A)^+^ RNA purification were performed for all RNA samples using the Dynabeads^®^ mRNA DIRECT^™^ Micro Kit (Invitrogen^™^/Thermo Fisher Scientific, Loughborough, UK) according to the manufacturer’s instructions. The purified poly(A)^+^ RNA was then used to generate strand-specific RNA-seq libraries using the ScriptSeq^™^ v2 RNA-Seq Library Preparation Kit, the ScriptSeq^™^ Index PCR Primers (Sets 1 to 4) and the FailSafe^™^ PCR enzyme system (all sourced from Epicentre^®^/Illumina^®^ Inc., Madison, WI, USA) according to the manufacturer’s instructions. RNA-seq libraries were purified using the Agencourt^®^ AMPure^®^ XP system (Beckman Coulter Genomics, Danvers, MA, USA) according to the manufacturer’s instructions for double size selection (0.75× followed by 1.0× ratio).

RNA-seq libraries were quantified using a Qubit^®^ fluorometer and Qubit^®^ dsDNA HS Assay Kit (Invitrogen^™^/Thermo Fisher Scientific), while library quality checks were performed using an Agilent 2100 Bioanalyzer and High Sensitivity DNA Kit (Agilent Technologies Ltd.). Individually barcoded RNA-seq libraries were pooled in equimolar quantities and the quantity and quality of the final pooled libraries (two different pools in total) were assessed as described above. RNA-seq library sample pool construction and barcode index sequences used to uniquely tag RNA-seq libraries in each pool are also detailed in Supplementary Table 1.

Cluster generation and high-throughput sequencing of the pooled RNA-seq libraries were performed using an Illumina^®^ HiSeq^™^ 2000 Sequencing System at the MSU Research Technology Support Facility (RTSF) Genomics Core (https://rtsf.natsci.msu.edu/genomics; Michigan State University, MI, USA). Each of the two pooled libraries were sequenced independently on five lanes split across multiple Illumina^®^ flow cells. The pooled libraries were sequenced as paired-end 2 × 100 nucleotide reads using Illumina^®^ version 5.0 sequencing kits. All RNA-seq data generated for this study have been deposited in the European Nucleotide Archive (ENA) database with project accession number **PRJEB44568**.

### 2.7. Bioinformatics and statistical analyses of RNA-seq data

Bioinformatics procedures were performed on a 32-core Linux Compute Server (4× AMD Opteron™ 6220 processors at 3.0 GHz with 8 cores each), with 256 GB of RAM, 24 TB of hard disk drive storage, and with Ubuntu Linux OS (version 14.04.4 LTS). All of the bioinformatics and statistical workflow scripts (bash, perl, and R programming languages) used are available from a public GitHub repository (https://github.com/UCD-Animal-Genomics-Group/PPD-b_RNA-sequencing) and **Fig. 2B** shows the RNA-seq bioinformatics and statistical workflow. Deconvolution was performed by the MSU RTSF Genomics Core using a pipeline that simultaneously demultiplexed and converted pooled sequence reads into discrete FASTQ files for each RNA-seq sample with no barcode index mismatches permitted.

The RNA-seq FASTQ sequence read data for each of the 80 samples were then downloaded from the MSU RTSF Genomics Core FTP server and the quality of individual raw RNA-seq sample library files was assessed with the FastQC software (version 0.11.9) [37]. Using the ngsShoRT tool (version 2.2) [38], the filtering strategy consisted of: (1) removal of paired-end reads containing adapter sequence contamination (allowing up to three mismatches); (2) removal of paired-end reads of poor quality (i.e., at least one of the reads containing ≥ 25% of bases with a Phred quality score below 20); (3) removal of paired end reads that did not meet the required minimum length of 100 nucleotides. The quality of filtered FASTQ files was then re-evaluated using the FastQC software. Paired-end reads, from each filtered individual library, were aligned to the ARS-UCD1.2 *B. taurus* reference genome (Accession number GCF_002263795.1) [39] using the STAR aligner (version 2.7.8a) [40]. FASTQ files originating from the same library sample, which were sequenced over multiple lanes, were launched together in a single job within STAR. Following this, the aligned BAM files were quality-assessed with FastQC. For each library, raw counts for each gene were obtained using the featureCounts software from the Subread package (version 1.6.4) [41], with parameters set to unambiguously assign uniquely aligned paired-end reads in a stranded manner to the exons of genes within the NCBI *Bos taurus* Annotation Release 106 genomic GFF file (**GCF_002263795.1**). Statistical analyses were performed on a MacBook Pro with a dual-core 2.9 GHz Intel^®^ Core™ i5 processor, 16 GB RAM, 500 GB SSD storage, and with the Darwin OS (version 17.0 - macOS Big Sur 10.16).

Raw gene count outputs from featureCounts were used to perform differential gene expression analysis within an R-based workflow (within the RStudio IDE version 1.4.1106 [42], running R version 4.1.0 [43]) with the edgeR package (version 3.34.0) [44] from the Bioconductor project (version 3.13) [45]. Data wrangling was performed with the tidyverse R package collection (version 1.3.1) [46], dplyr (version 1.0.7) [47], magrittr (version 2.0.1) [48], and biobroom (version 1.24.0) [49] R packages. Charts were generated with the following packages: ggplot2 version 3.3.5 [50], Cairo version 1.5-12.2 [51], ComplexHeatmap version 2.9.3 [52], cowplot version 1.1.1 [53], ggfortify version 0.4.12 [54], ggrepel version 0.9.1 [55], and ggridges version 0.5.3 [56].

Normalisation factors for each library were calculated using the trimmed mean of M-values method (TMM) [57]. Differentially expressed (DE) genes between the PPD-b-stimulated animal group and the control non-stimulated group at each time point (-1 wk pre-infection; +1 wk, +2 wk, and +10 wk post-infection) were identified using a paired-sample design with animal as a blocking factor. Differential expression was evaluated by fitting a quasi-likelihood (QL) negative binomial (NB) generalized linear model (GLM) for each gene [58, 59], while also conducting an empirical Bayes (EB) moderation of the genewise QL dispersions [60]; and applying genewise statistical tests with EB QL F-tests [58]. Correction for multiple testing was done via the Benjamini-Hochberg method (B-H) [61] with a false discovery rate (FDR) adjusted *P*-value threshold of < 0.001 for genes considered to be DE.

### 2.8. Ingenuity Pathway Analysis of RNA-seq gene expression

The Ingenuity^®^ Pathway Analysis (IPA) software package [62] with the Ingenuity^®^ Knowledge Base (Qiagen, Redwood City, CA, USA; release date December 2021) was used to identify overrepresented (enriched) canonical pathways for sets of DE genes at each post-infection time point. For each timepoint, IPA^®^ Core Analysis was performed using the default settings with the user gene expression data set as the background, high predicted confidence and all nodes selected. A stringent Benjamini–Hochberg (B-H) FDR [61] threshold of < 0.001 was applied to filter the input DE genes and generate gene lists of a suitable size for the IPA^®^ Core Analysis. For identification of overrepresented canonical pathways, a B-H *P*-value adjustment was also applied with an FDR *P*_adj._ threshold < 0.05.

## 3. Results

### 3.1. RNA-seq summary statistics and exploratory data analysis

Filtering of sequence reads to remove adapter-dimer contamination and low-quality reads yielded a mean of 17,870,217 ± 2,521,780 reads per individual sample library (*n* = 80 libraries and ± SD); this represented a mean of 82% of reads that passed this step. Filtered reads were then aligned to the ARS-UCD1.2 *B. taurus* genome build, with a mean of 15,567,036 ± 2,054,294 reads (87.3%) that uniquely mapped to the genome, 547,884 ± 178,455 reads (3.08%) that mapped to multiple genomic locations, 22,241 ± 6,747 reads (0.12%) mapped to excessive numbers of loci, and 1,733,056 ± 625,662 reads (9.5%) that did not map to any genomic location. The mean mapped length was 197.1 ± 0.5 bp. Individual sample information is provided in Supplementary Table 2 and **Fig. 3** details the RNA-seq summary statistics.

**Fig. 3:**
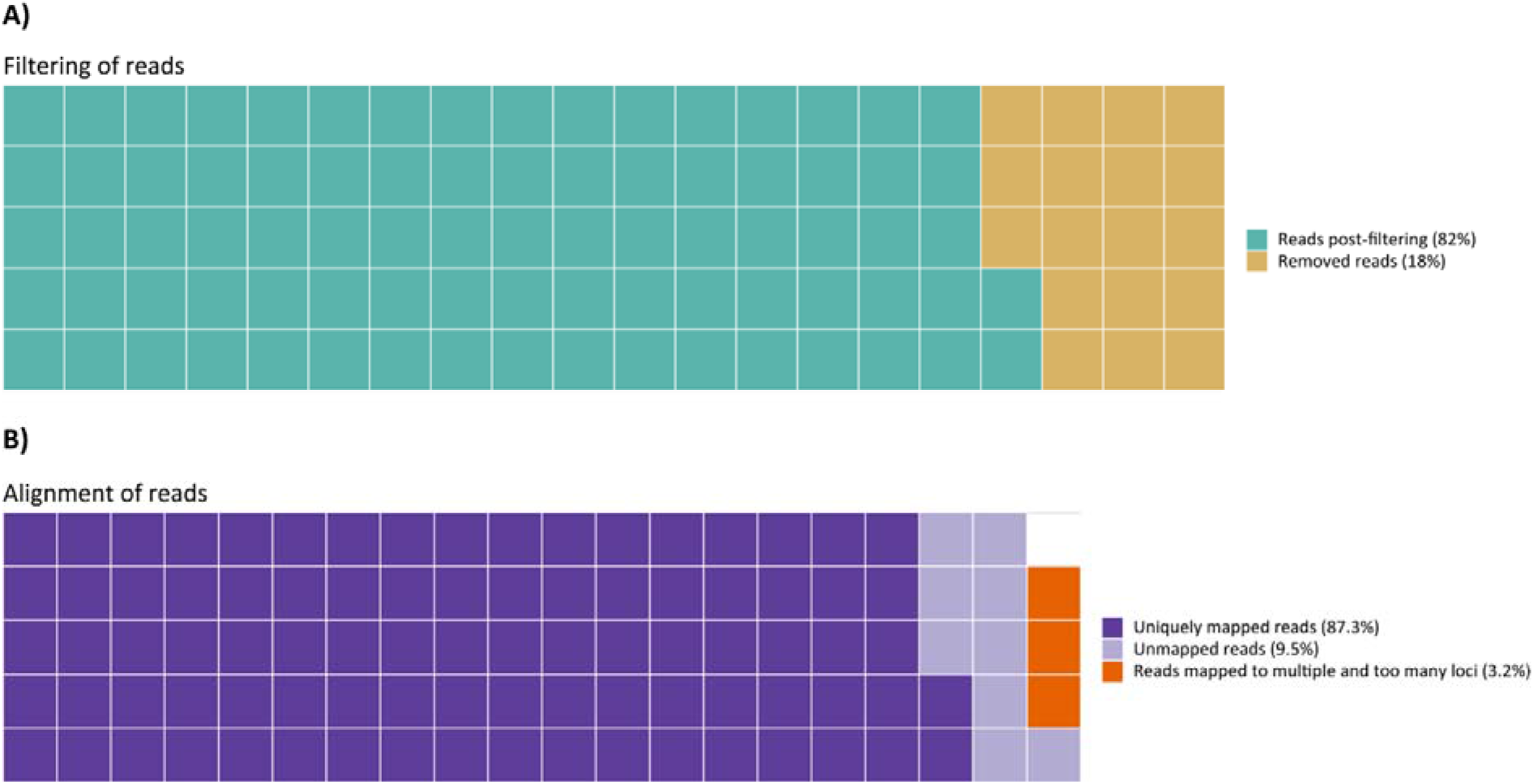
Waffle charts of RNA-seq summary statistics. **A.** Mean percentage of sequence reads per library that were retained after filtering of reads; and those removed due to adapter sequence contamination or low sequence quality. **B.** Mean percentage of sequence reads per library that aligned to unique locations in the *B. taurus* reference genome, multiple locations and too many loci, and reads that did not align to any genomic locations.

Supplementary Figures 1 and 2 show that none of the libraries exhibited an abnormal distribution before or after filtering of lowly expressed genes. Removal of lowly expressed genes prior to differential expression analysis has been shown to improve power of detection and reduce the number of false positive errors [63, 64]. Expression data from filtered genes was then used for Principal Component Analysis (PCA) to visualise differences among the expression profiles of individual libraries at each time point for both PPD-b-stimulated and control non-stimulated groups (Supplementary Figure 3).

### 3.2. Differential gene expression of peripheral blood samples in response to PPD-b stimulation

Statistical analysis of the RNA-seq gene expression data using the edgeR package with a B-H FDR-adjusted *P*-value threshold < 0.001 revealed 374 differentially expressed (DE) genes in PPD-b-stimulated -1 wk pre-infection versus the control non-stimulated -1 wk pre-infection group; 280 of these genes showed increased expression, while 94 exhibited decreased expression. Similarly, 349 DE genes (FDR *P*_adj._ < 0.001) were observed for PPD-b-stimulated +1 wk post-infection: 203 displayed increased expression and 146 showed decreased expression relative to the control non-stimulated +1 wk post-infection group. The PPD-b-stimulated +2 wk post-infection group, exhibited 1,616 DE genes (FDR *P*_adj._ < 0.001), with increased expression for 902 genes and decreased expression for 714 genes relative to the control non-stimulated +2 wk post-infection group. Finally, the largest number of DE genes (FDR *P*_adj._ < 0.001) were observed for PPD-b-stimulated +10 wk post-infection where 5,590 DE genes were detected, and 2,759 genes exhibited increased expression and 2,831 genes showed decreased expression relative to the control non-stimulated +10 wk post-infection group. **Fig. 4A** summarises these results and **Fig. 4B** shows intersecting DE genes across all comparisons.

**Fig. 4:**
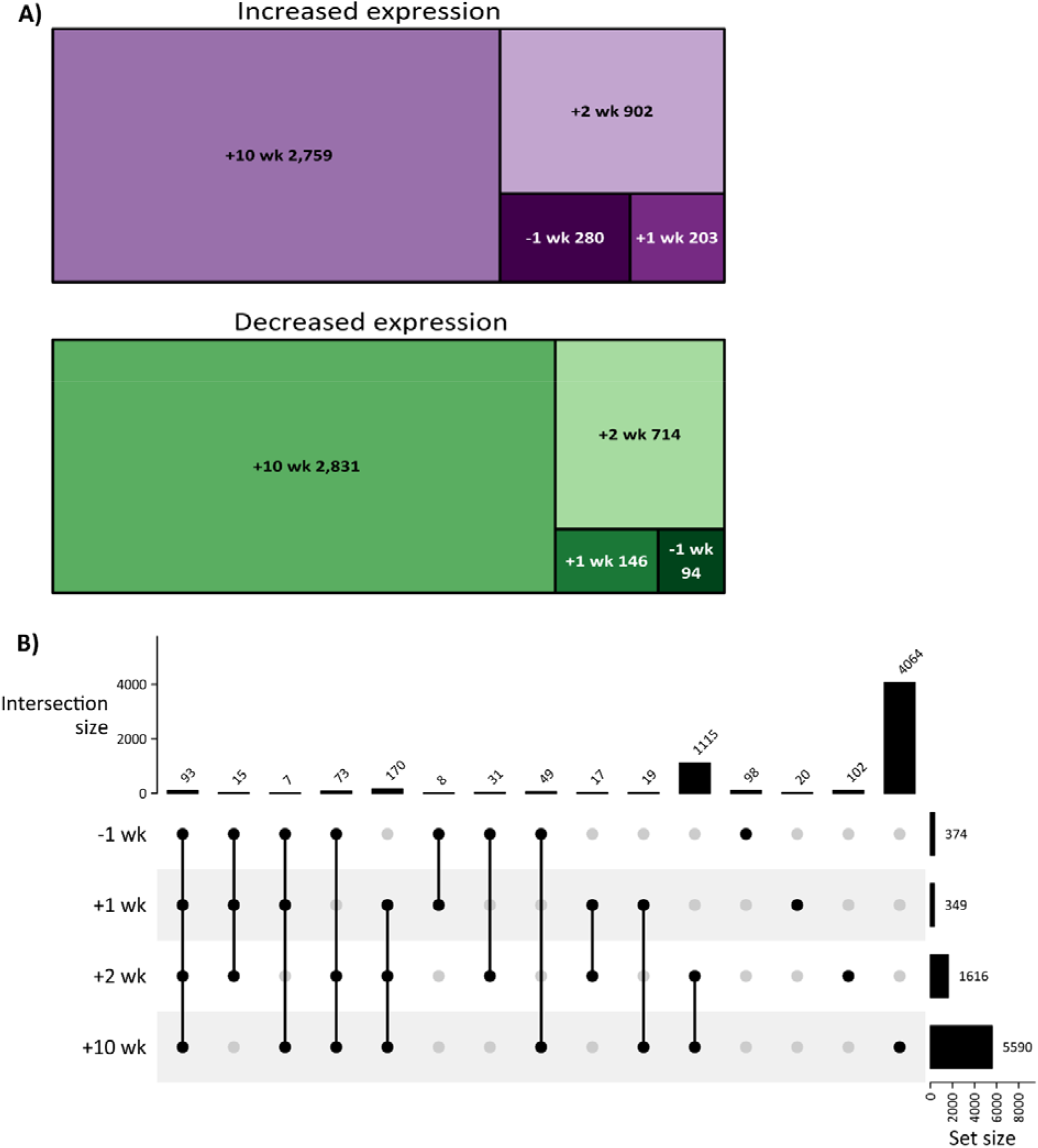
Statistically significant differentially expressed genes. **A.** Treemaps showing either increased expression (purple) or decreased expression (green) at each PPD-b-stimulated time point (-1 wk pre-infection; +1 wk, +2 wk, and +10 wk post infection) relative to the non-stimulated controls. **B.** UpSet plot displaying DE gene intersections among all comparisons. The threshold for statistical significance of differentially expressed genes was a Benjamini-Hochberg FDR-adjusted *P* value < 0.001.

**Tables 1–4** provide information on the top twenty DE genes that exhibited increased and decreased expression at each of the four time point comparisons. Among the top DE genes identified (**Tables 1–4**), higher expression of genes encoding cytokine receptors (*IL3RA*, -1 wk), interleukins (*IL17A* +10 wk), and genes induced by interferon (*GBP5*, *GBP4*, *LOC512486* – *GBP1*, +2 wk; *CXCL10*, +10 wk; *LOC100336669* – *GBP4*, *LOC511531* – another *GBP1* ortholog, +2 wk and + 10 wk) was observed. On the other hand, lower expression was observed for genes encoding proteins involved in antimicrobial activity (*RNASE6*, +1 wk; *RNASE4* +2 wk), C-type lectin receptors (*CLEC4A*, -1 wk; *CLEC7A*, +1 wk; *CLEC12A*, +10 wk), and inhibition of signal transduction (*ARRB2*, +10 wk). Detailed information on DE genes for each contrast between PPD-b-stimulated and control non-stimulated time points are provided in Supplementary Tables 3–6, while Supplementary Table 7 provides additional information on the DE genes that were common to all comparisons (93 genes, **Figure 4B**).

**Table 1:**
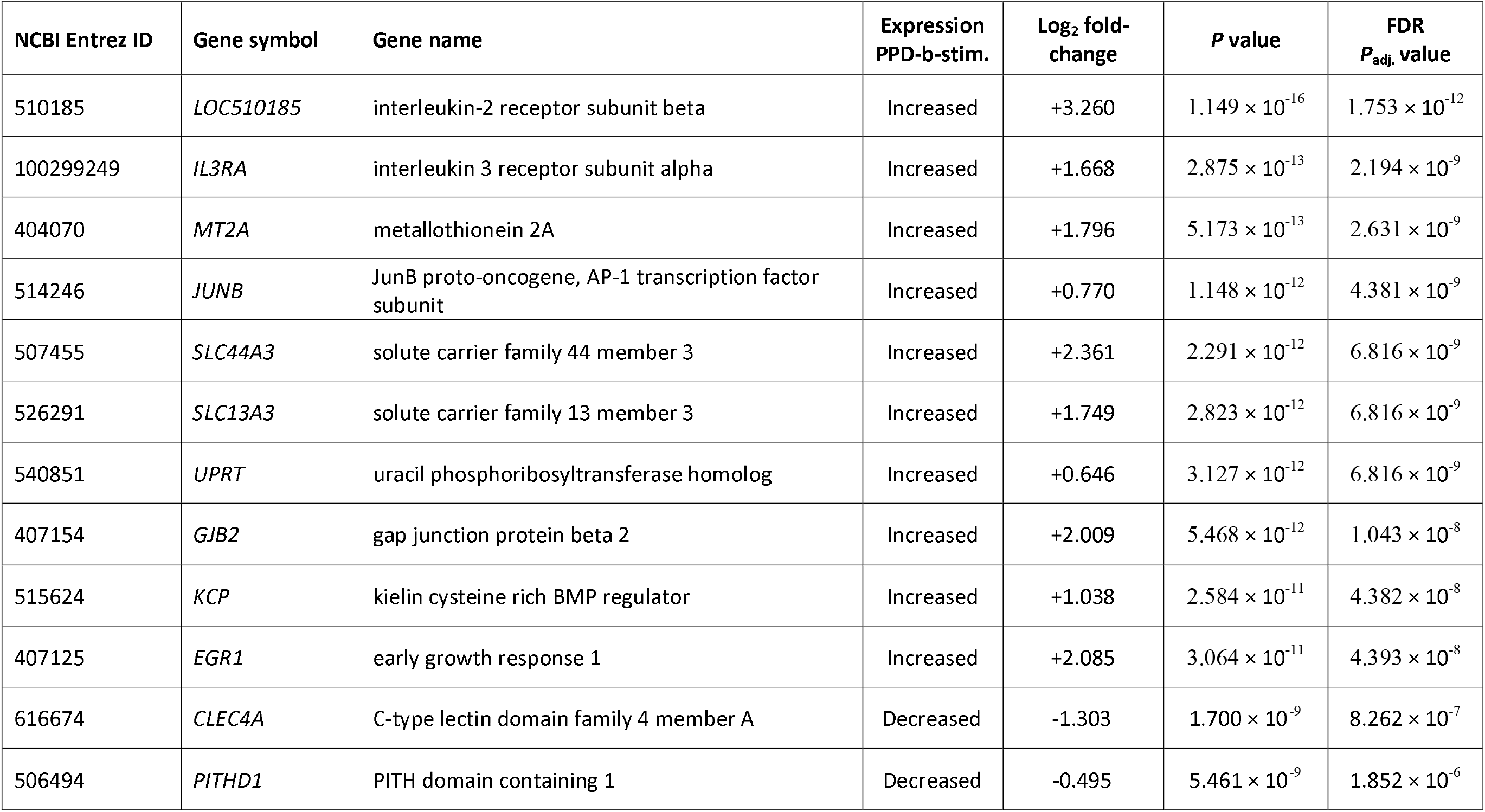
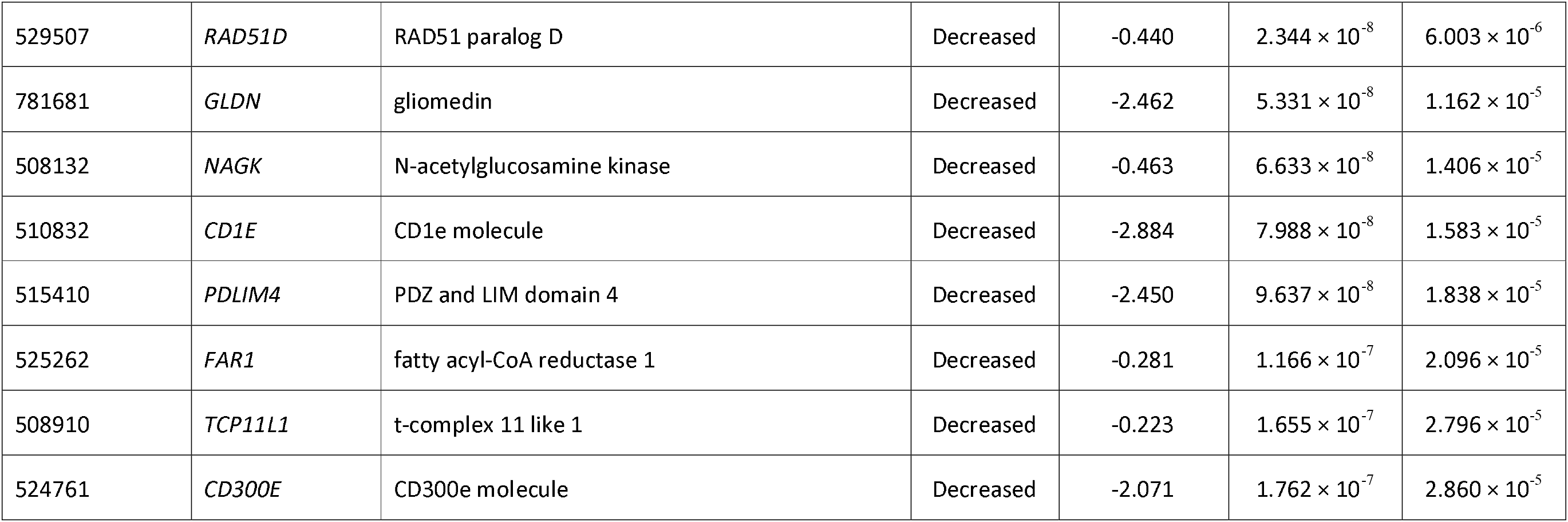
Top ten differentially expressed genes exhibiting increased and decreased expression in the PPD-b-stimulated -1 wk pre-infection group relative to the control non-stimulated -1 wk pre-infection group. Genes are ranked by statistical significance.

**Table 2:** Top ten differentially expressed genes exhibiting increased and decreased expression in the PPD-b-stimulated +1 wk post-infection group relative to the control non-stimulated +1 wk post-infection group. Genes are ranked by statistical significance.

| NCBI Entrez ID | Gene symbol | Gene name | Expression PPD-b-stim. | Log<sub>2</sub> fold-change | P value | FDR P<sub>adj.</sub> value |
| --- | --- | --- | --- | --- | --- | --- |
| 510185 | *LOC510185* | interleukin-2 receptor subunit beta | Increased | +3.060 | $2.017 \times 10^{-15}$ | $3.077 \times 10^{-11}$ |
| 281309 | *MMP3* | matrix metalloproteinase 3 | Increased | +3.895 | $3.276 \times 10^{-13}$ | $2.499 \times 10^{-9}$ |
| 526291 | *SLC13A3* | solute carrier family 13 member 3 | Increased | +1.812 | $4.958 \times 10^{-13}$ | $2.522 \times 10^{-9}$ |
| 281044 | *CCL8* | chemokine (C-C motif) ligand 8 | Increased | +3.025 | $3.077 \times 10^{-12}$ | $1.174 \times 10^{-8}$ |
| 280826 | *IL6* | interleukin 6 | Increased | +2.416 | $2.763 \times 10^{-10}$ | $6.504 \times 10^{-7}$ |
| 782652 | *FCGR2A* | Fc fragment of IgG receptor IIa | Increased | +2.027 | $2.984 \times 10^{-10}$ | $6.504 \times 10^{-7}$ |
| 282201 | *CRABP1* | cellular retinoic acid binding protein 1 | Increased | +2.313 | $5.704 \times 10^{-10}$ | $1.088 \times 10^{-6}$ |
| 614655 | *DESI1* | desumoylating isopeptidase 1 | Increased | +0.546 | $6.502 \times 10^{-10}$ | $1.102 \times 10^{-6}$ |
| 112441463 | *LOC112441463* | interstitial collagenase-like | Increased | +3.462 | $9.409 \times 10^{-10}$ | $1.436 \times 10^{-6}$ |
| 526981 | *MMP12* | matrix metalloproteinase 12 | Increased | +2.980 | $1.538 \times 10^{-10}$ | $1.473 \times 10^{-6}$ |
| 506494 | *PITHD1* | PITH domain containing 1 | Decreased | -0.549 | $2.863 \times 10^{-10}$ | $6.504 \times 10^{-7}$ |
| 532865 | *CLEC7A* | C-type lectin domain containing 7A | Decreased | -2.514 | $2.880 \times 10^{-9}$ | $2.197 \times 10^{-6}$ |
| 112448804 | *LOC112448804* | uncharacterized LOC112448804 | Decreased | -3.729 | $1.660 \times 10^{-8}$ | $8.444 \times 10^{-6}$ |
| 282341 | *RNASE6* | ribonuclease A family member k6 | Decreased | -1.509 | $2.739 \times 10^{-8}$ | $1.267 \times 10^{-5}$ |
| 511960 | *CYP27A1* | cytochrome P450, family 27, subfamily A, polypeptide 1 | Decreased | -2.843 | $4.432 \times 10^{-8}$ | $1.879 \times 10^{-5}$ |
| 508888 | *PLCB2* | phospholipase C beta 2 | Decreased | -0.414 | $7.540 \times 10^{-8}$ | $2.739 \times 10^{-5}$ |
| 524761 | *CD300E* | CD300e molecule | Decreased | -2.115 | $8.496 \times 10^{-8}$ | $2.886 \times 10^{-5}$ |
| 516258 | *TRMT10B* | tRNA methyltransferase 10B | Decreased | -0.287 | $1.114 \times 10^{-7}$ | $3.541 \times 10^{-5}$ |
| 282118 | *VCAM1* | vascular cell adhesion molecule 1 | Decreased | -3.068 | $1.176 \times 10^{-7}$ | $3.625 \times 10^{-5}$ |
| 533151 | *ABCC3* | ATP binding cassette subfamily C member 3 | Decreased | -0.984 | $1.328 \times 10^{-7}$ | $3.790 \times 10^{-5}$ |

**Table 3:** Top ten differentially expressed genes exhibiting increased and decreased expression in the PPD-b-stimulated +2 wk post-infection group relative to the control non-stimulated +2 wk post-infection group. Genes are ranked by statistical significance.

| NCBI Entrez ID | Gene symbol | Gene name | Expression PPD-b-stim. | Log<sub>2</sub> fold-change | P value | FDR P<sub>adj.</sub> value |
| --- | --- | --- | --- | --- | --- | --- |
| 516949 | *GBP5* | guanylate binding protein 5 | Increased | +2.740 | $1.302 \times 10^{-22}$ | $1.987 \times 10^{-18}$ |
| 613313 | *GBP4* | guanylate binding protein 4 | Increased | +2.407 | $5.507 \times 10^{-22}$ | $3.148 \times 10^{-18}$ |
| 512486 | *LOC512486* | interferon-induced guanylate-binding protein 1 | Increased | +2.344 | $6.190 \times 10^{-22}$ | $3.148 \times 10^{-18}$ |
| 510814 | *STAT1* | signal transducer and activator of transcription 1 | Increased | +1.732 | $8.714 \times 10^{-22}$ | $3.324 \times 10^{-18}$ |
| 510185 | *LOC510185* | interleukin-2 receptor subunit beta | Increased | +4.098 | $1.116 \times 10^{-21}$ | $3.405 \times 10^{-18}$ |
| 100336669 | *LOC100336669* | guanylate-binding protein 4 | Increased | +3.233 | $1.412 \times 10^{-20}$ | $3.590 \times 10^{-17}$ |
| 511531 | *LOC511531* | guanylate-binding protein 1 | Increased | +2.029 | $2.312 \times 10^{-20}$ | $5.039 \times 10^{-17}$ |
| 786372 | *LOC786372* | golgin subfamily A member 6-like protein 22 | Increased | +2.368 | $6.580 \times 10^{-20}$ | $1.255 \times 10^{-16}$ |
| 533834 | *CD274* | CD274 molecule | Increased | +2.218 | $1.057 \times 10^{-19}$ | $1.792 \times 10^{-16}$ |
| 789216 | *IRF1* | interferon regulatory factor 1 | Increased | +2.016 | $2.727 \times 10^{-19}$ | $4.160 \times 10^{-16}$ |
| 516545 | *FAM210B* | family with sequence similarity 210 member B | Decreased | -0.865 | $2.105 \times 10^{-11}$ | $2.170 \times 10^{-9}$ |
| 618814 | *HACD4* | 3-hydroxyacyl-CoA dehydratase 4 | Decreased | -2.245 | $1.304 \times 10^{-10}$ | $1.053 \times 10^{-8}$ |
| 511755 | *SFXN3* | sideroflexin 3 | Decreased | -0.993 | $3.303 \times 10^{-10}$ | $2.434 \times 10^{-8}$ |
| 616089 | *RNASE4* | ribonuclease A family member 4 | Decreased | -2.443 | $6.563 \times 10^{-10}$ | $4.573 \times 10^{-8}$ |
| 511960 | *CYP27A1* | cytochrome P450, family 27, subfamily A, polypeptide 1 | Decreased | -3.536 | $7.997 \times 10^{-10}$ | $5.471 \times 10^{-8}$ |
| 514877 | *RASGEF1A* | RasGEF domain family member 1A | Decreased | -1.937 | $8.123 \times 10^{-10}$ | $5.533 \times 10^{-8}$ |
| 535260 | *FAM151B* | family with sequence similarity 151 member B | Decreased | -0.784 | $8.401 \times 10^{-10}$ | $5.697 \times 10^{-8}$ |
| 282879 | *HGF* | hepatocyte growth factor | Decreased | -2.378 | $8.536 \times 10^{-10}$ | $5.763 \times 10^{-8}$ |
| 510832 | *CD1E* | CD1e molecule | Decreased | -3.763 | $1.013 \times 10^{-9}$ | $6.679 \times 10^{-8}$ |
| 112448804 | *LOC112448804* | uncharacterized LOC112448804 | Decreased | -4.890 | $1.084 \times 10^{-9}$ | $7.051 \times 10^{-8}$ |

**Table 4:** Top ten differentially expressed genes exhibiting increased and decreased expression in the PPD-b-stimulated +10 wk post-infection group relative to the control non-stimulated +10 wk post-infection group. Genes are ranked by statistical significance.

| NCBI Entrez ID | Gene symbol | Gene name | Expression PPD-b-stim. | Log<sub>2</sub> fold-change | P value | FDR P<sub>adj.</sub> value |
| --- | --- | --- | --- | --- | --- | --- |
| 614655 | *DESI1* | desumoylating isopeptidase 1 | Increased | +1.723 | $7.800 \times 10^{-36}$ | $6.857 \times 10^{-32}$ |
| 506030 | *IL17F* | interleukin 17F | Increased | +7.606 | $8.988 \times 10^{-36}$ | $6.857 \times 10^{-32}$ |
| 281237 | *IFNG* | interferon gamma | Increased | +7.848 | $9.931 \times 10^{-35}$ | $5.051 \times 10^{-31}$ |
| 280701 | *PPA1* | inorganic pyrophosphatase 1 | Increased | +2.620 | $1.644 \times 10^{-34}$ | $6.270 \times 10^{-31}$ |
| 338034 | *MYBL1* | MYB proto-oncogene like 1 | Increased | +2.724 | $5.534 \times 10^{-34}$ | $1.689 \times 10^{-30}$ |
| 511531 | *LOC511531* | guanylate-binding protein 1 | Increased | +3.208 | $3.570 \times 10^{-33}$ | $9.078 \times 10^{-30}$ |
| 100336669 | *LOC100336669* | guanylate-binding protein 4 | Increased | +4.802 | $7.933 \times 10^{-33}$ | $1.729 \times 10^{-29}$ |
| 104971647 | *LOC104971647* | uncharacterized LOC104971647 | Increased | +3.327 | $2.440 \times 10^{-32}$ | $4.654 \times 10^{-29}$ |
| 615107 | *CXCL10* | C-X-C motif chemokine ligand 10 | Increased | +6.322 | $2.760 \times 10^{-32}$ | $4.679 \times 10^{-29}$ |
| 282863 | *IL17A* | interleukin 17A | Increased | +7.456 | $4.233 \times 10^{-32}$ | $6.459 \times 10^{-29}$ |
| 281638 | *ARRB2* | arrestin beta 2 | Decreased | -1.092 | $3.071 \times 10^{-23}$ | $4.184 \times 10^{-21}$ |
| 504317 | *LRP4* | LDL receptor related protein 4 | Decreased | -2.379 | $2.102 \times 10^{-22}$ | $2.486 \times 10^{-20}$ |
| 101902671 | *XG* | Xg glycoprotein | Decreased | -1.913 | $2.200 \times 10^{-21}$ | $2.268 \times 10^{-19}$ |
| 540667 | *MARCH8* | membrane associated ring-CH-type finger 8 | Decreased | -0.568 | $7.174 \times 10^{-20}$ | $5.731 \times 10^{-18}$ |
| 515399 | *STAG3* | stromal antigen 3 | Decreased | -1.799 | $2.401 \times 10^{-19}$ | $1.761 \times 10^{-17}$ |
| 511220 | *ARHGEF11* | Rho guanine nucleotide exchange factor 11 | Decreased | -2.270 | $5.174 \times 10^{-19}$ | $3.621 \times 10^{-17}$ |
| 516875 | *IPCEF1* | interaction protein for cytohesin exchange factors 1 | Decreased | -0.955 | $5.478 \times 10^{-19}$ | $3.817 \times 10^{-17}$ |
| 509416 | *WC1-12* | WC1 isolate DV10 | Decreased | -1.439 | $7.043 \times 10^{-19}$ | $4.798 \times 10^{-17}$ |
| 524183 | *TMEM164* | transmembrane protein 164 | Decreased | -0.749 | $9.037 \times 10^{-19}$ | $5.995 \times 10^{-17}$ |
| 508740 | *CLEC12A* | C-type lectin domain family 12 member A | Decreased | -2.770 | $9.677 \times 10^{-19}$ | $6.364 \times 10^{-17}$ |

A general response to PPD-b stimulation was observed that was composed of 93 genes that were consistently differentially expressed across all time points, irrespective of infection status (Supplementary Table 7). Among these, decreased expression was observed for genes encoding ATP-binding cassette (ABC) transporter (*ABCC3*), and transmembrane (*CD1B*, *CD1E*) and cell surface (*CD59*, *CD300E*) glycoproteins. In addition, genes encoding chemokines (*CCL17*, *CCL8*, *XCL2*, *GRO1*), interleukins and receptor subunits (*IL6*, *IL27*, *LOC510185* – *IL2RB*), matrix metallopeptidase (*MMP3*), and colony stimulating factor 3 cytokine (*CSF3*) were increased in expression across all time points. Detection of this general inflammatory response indicates that expression changes for these genes might not be useful to distinguish between *M. bovis*-infected and non-infected animals after PPD-b stimulation.

### 3.3. Pathway analysis

The numbers of genes used for IPA^®^ Core Analysis are shown in Supplementary Table 9. The top five canonical pathways for each comparison are shown in **Table 5**. In addition, the top 10 canonical pathways for each time point comparison are shown as bar charts in Supplementary Figures 4–7. The IPA^®^ canonical pathway that was ranked first at the -1 wk pre-infection time point according to overrepresentation of DE genes is the *Coronavirus Pathogenesis Pathway* (**Table 5**), which has recently been incorporated into the Ingenuity^®^ Knowledge Base [65, 66]. Nineteen of the 170 genes (11.2%) in this pathway were differentially expressed at the -1 wk pre-infection time point. As shown in **Table 5**, the other top-ranked pathways for the -1 wk and the three post-infection time points (+1, +2 and +10 wk) include immune system and signalling pathways.

**Table 5:** Top five overrepresented IPA^®^ canonical pathways for each of the four time point comparisons between PPD-b stimulated and unstimulated peripheral blood samples.

| Time point | Top canonical pathways | *P*-value | w | % genes overlapping pathway | No. of genes overlapping pathway |
| --- | --- | --- | --- | --- | --- |
| -1 wk | Coronavirus Pathogenesis Pathway | $2.14 \times 10^{-7}$ | 4.04 | 11.2% | 19/170 |
| | iNOS Signalling | $2.45 \times 10^{-6}$ | 3.29 | 20.5% | 9/44 |
| | IL-17 Signalling | $1.11 \times 10^{-5}$ | 2.81 | 10.9% | 14/128 |
| | Tumour Microenvironment Pathway | $4.62 \times 10^{-5}$ | 2.31 | 10.2% | 13/127 |
| | Granulocyte Adhesion and Diapedesis | $6.96 \times 10^{-5}$ | 2.23 | 11.3% | 11/97 |
| +1 wk | Granulocyte Adhesion and Diapedesis | $2.43 \times 10^{-13}$ | 10 | 20.6% | 20/97 |
| | Agranulocyte Adhesion and Diapedesis | $4.37 \times 10^{-13}$ | 10 | 18.8% | 21/112 |
| | Role of Cytokines in Mediating Communication between Immune Cells | $1.96 \times 10^{-10}$ | 7.54 | 40.0% | 10/25 |
| | Differential Regulation of Cytokine Production in Macrophages and T Helper Cells by IL-17A and IL-17F | $8.68 \times 10^{-9}$ | 6.02 | 53.8% | 7/13 |
| | Airway Pathology in Chronic Obstructive Pulmonary Disease | $1.91 \times 10^{-8}$ | 5.77 | 20.3% | 12/59 |
| +2 wk | Granulocyte Adhesion and Diapedesis | $5.33 \times 10^{-15}$ | 11.5 | 42.3% | 41/97 |
| | Role of Hypercytokinemia/hyperchemokineemia in the Pathogenesis of Influenza | $3.14 \times 10^{-14}$ | 11 | 54.9% | 28/51 |
| | Agranulocyte Adhesion and Diapedesis | $3.48 \times 10^{-13}$ | 10.2 | 37.5% | 42/112 |
| | Role of Cytokines in Mediating Communication between Immune Cells | $1.67 \times 10^{-12}$ | 9.62 | 72.0% | 18/25 |
| | Pyroptosis Signalling Pathway | $1.63 \times 10^{-11}$ | 8.72 | 43.9% | 29/66 |
| +10 wk | Crosstalk between Dendritic Cells and Natural Killer Cells | $2.19 \times 10^{-8}$ | 5.1 | 73.8% | 45/61 |
| | Hepatic Cholestasis | $2.90 \times 10^{-8}$ | 5.1 | 62.1% | 82/132 |
| | Granulocyte Adhesion and Diapedesis | $3.53 \times 10^{-8}$ | 5.1 | 66.0% | 64/97 |
| | FAT10 Signalling Pathway | $1.78 \times 10^{-7}$ | 4.52 | 74.5% | 38/51 |
| | Cardiac Hypertrophy Signalling (Enhanced) | $2.64 \times 10^{-7}$ | 4.48 | 50.7% | 204/402 |

In cases where the same canonical pathway is overrepresented across multiple time points there is increased perturbation of the pathway as the infection progresses: *Granulocyte Adhesion and Diapedesis* at -1, +1, +2 and +10 wk post-infection, and *Agranulocyte Adhesion and Diapedesis* and *Role of Cytokines in Mediating Communication between Immune Cells* at +1 and +2 wk post-infection. This is particularly apparent in the case of the *Granulocyte Adhesion and Diapedesis* pathway, which is the top-ranked pathway for the first two post-infection time points (**Table 5**). At the -1 wk time point 11 of the 97 genes in the pathway (11.3%) are differentially expressed. This increases to 20 genes (20.6%) for the +1 wk time point, 41 genes (42.3%) for the +2 wk time point and 64 genes (66.0%) for the +10 wk time point. Pathway diagrams for the *Granulocyte Adhesion and Diapedesis* pathway at each time point are shown in Supplementary Figures 8–11.

## 4. Discussion

Mycobacterial infections, including *M. bovis* infection of cattle that causes bTB disease, generate a spectrum of host immune responses, involving a complex interplay between both innate and adaptive immune mechanisms [67–69]. However, it is well established that peripheral immune responses of cattle infected with *M. bovis* can provide information about the responses at the site of active infection and disease [70]. Therefore, we have used RNA□seq transcriptomic profiling of bovine peripheral blood following an overnight in vitro incubation with PPD-b to evaluate transcriptional changes and identify blood transcriptional modules enriched after mycobacterial antigen (tuberculin – PPD-b) stimulation of samples from cattle across four time points (-1 wk pre-infection; +1 wk, +2 wk, and +10 wk post-infection).

### 4.1. PPD-b induced perturbation of the PBL transcriptome from *M. bovis*-infected cattle

As shown in **Figure 4**, differential gene expression was observed in all four PPD-b-stimulated sample groups compared to parallel non-stimulated controls at the same time points (-1 wk pre-infection; +1 wk, +2 wk, and +10 wk post-infection). Our group has previously analysed RNA-seq data from non-stimulated peripheral blood samples from the same cohort of naïve animals across an expanded infection time course [29]. In that study, results from each of five post-infection time points (+1 wk, +2 wk, +6 wk, +10 wk and +12 wk) versus one control pre-infection time point (-1 wk) revealed smaller numbers of DE genes at +1 wk post-infection (37 genes with increased expression and 20 with decreased expression), +2 wk post-infection (83 increased and 10 decreased), and +10 wk post-infection (1,278 increased and 1,305 decreased). Additional DE genes were detected at +6 wk post-infection (415 increased and 272 decreased), and +12 wk post-infection (222 increased and 116 decreased) [29]. It is also important to note that a less stringent statistical threshold (FDR *P*_adj._ < 0.05) was used to detect DE genes in this parallel study of non-stimulated peripheral blood samples.

The present study, therefore, demonstrates that in vitro PPD-b stimulation of non-infected and *M. bovis*-infected peripheral blood elicits a more substantial perturbation of the host transcriptome, even though a very stringent B-H FDR-adjusted statistical threshold was used (FDR *P*_adj._ < 0.001). Similar to the experiment described in the present study—RNA-seq gene expression using whole blood samples stimulated overnight at 37°C with PPD-b versus a control non-stimulated group—a previous study using a different cohort of cattle has reported a large number of DE genes (|log_2_FC| >1 cut-off; B-H FDR *P*_adj._ < 0.05 threshold) at +2 wk (day 14: 2,622 genes), and +6 wk (day 42: 1,446 genes) post-infection with 1.12 × 10^4^ CFU of *M. bovis* AF2122/97 [71]. Although the same pathogenic strain of *M. bovis* and inoculum delivery route was used for the experimental infection described in the current study, Villarreal-Ramos et al. used a 5.6-fold higher infective dose [71]. Additional differences between the two studies include the cattle breed (a Limousin × Simmental cross population used by Villarreal-Ramos et al. versus Holstein-Friesian animals in the present study); the sex of the animals used (exclusively female versus exclusively male); and fold-change cut-off and statistical thresholds used for the detection of DE genes (absolute log_2_FC ≥ 1 and B-H FDR *P*_adj._ < 0.05, versus no fold-change cut-off and B-H FDR *P*_adj._ < 0.001).

In the present study, some genes exhibiting increased expression in bovine peripheral blood stimulated with PPD-b such as *CD274* (at -1, +1, +2 and +10 wk), *P2RY14* (at +2 wk and +10 wk), *S100A8* (at +10 wk), and *CALHM6* (at +2 and +10 wk), have also been shown to exhibit increased expression in human TB cases when compared to healthy controls [72]. The *FBXL5* gene was also overexpressed in human TB patients [72]; however, it was decreased in expression at +10 wk in the present study (Supplementary Table 6). Moreover, these genes were components of a panel of 12 top classifier genes for a diagnostic point-of-care test for human TB [72]. In addition, comparison of the results obtained in the present study demonstrated that statistically significant DE genes across the infection time course of PPD-b-stimulated peripheral bovine blood samples were also components of a 30-gene expression signature recently identified for clinical human TB [73]. These were *BATF2* and *CD274* (-1, +1, +2 and +10 wk); *C1QA*, *CALHM6*—aka *FAM26F*, and *PDCD1LG2* (+2 and +10 wk); and *C1QC*, *C1QB*, *CFB*, *FAM20A*, *GBP6*, and *VWA3B* (+10 wk). This human study also identified a 30-gene expression signature for subclinical human TB, and genes that overlapped the results obtained in the present study were *BATF2* (-1, +1, +2 and +10 wk); *GBP5* (+1, +2 and +10 wk); *HTRA1* (-1 wk); *C1QC*, *GBP6*, *H4C14*, and *TICAM2* (+10 wk).

A recent transcriptomics study compared RNA-seq data from human peripheral blood mononuclear cells (PBMC) obtained from active human TB cases and non-infected controls [74]. Using a similar methodology to that employed for the present study, the human PBMC samples were stimulated with *M. tuberculosis* purified protein derivative (PPD) for four hours prior to RNA extraction. Comparison of these PBMC data with the results from the present study (using a less stringent statistical threshold of FDR *P*_adj._ < 0.01) showed that 11.2% (11/98) of the human PBMC DE genes overlapped with DE genes at the -1 wk contrast in cattle; 16.3% (16/98) overlapped with DE genes at the +1 wk contrast in cattle; 26.5% (26/98) overlapped with DE genes at the +2 wk contrast in cattle; and 48.0% (47/98) overlapped with DE genes at the +10 wk contrast in cattle. The DE human PBMC genes that overlapped with all four bovine pre- and post-infection time points were *CXCL10*, *EDN1*, *GBP4*, *IL1RN*, *MMP1*, *RNF19B*, *SLC37A2*, and *SOCS2*. Notable DE human PBMC genes that overlapped with DE genes at the three post-infection time points in cattle included *ALOX5*, *CXCL5*, *MAOA*, *PDK4*, and *STAT5A*. Finally, additional DE human PBMC genes that overlapped with the +2 wk and +10 wk post-infection time points in cattle included *AGAP1*, *EEPD1*, *IFNG*, *IL26*, *TIMP3*, *SLC6A9*, *SPIRE2*, and *STAB1*.

### 4.2. Leukocyte migration pathways and antigen stimulation of PBL from *M. bovis*-infected cattle

The increasing perturbation of *Granulocyte Adhesion and Diapedesis* pathway across the infection time course is detailed in Supplementary Figures 8–11. Perturbation of this pathway in response to *M. bovis* infection and subsequent stimulation of peripheral blood with PPD-b is likely a consequence of inflammatory responses that trigger extravasation of neutrophils and other leukocytes from the vasculature to infected tissues and adhesion to the extracellular matrix (ECM) [75, 76]. In this regard, a previous study of unstimulated PBL from *M. bovis*-infected and non-infected control cattle showed enrichment and suppression of the related IPA^®^ *Leukocyte Extravasation Signalling* pathway [30]. In addition, enrichment of a very similar KEGG pathway (*Leukocyte Transendothelial Migration*) was also observed in a recent RNA-seq-based transcriptomics study of peripheral blood mononuclear cells (PBMC) from control healthy cattle versus animals that were classified PCR-positive for bTB disease using an *M. bovis mpb70* gene nested PCR amplicon with nasal swabs [26].

### 4.3. Core responses to mycobacterial antigen stimulation that depend on infection status

Using peripheral blood and RT-qPCR, a four-gene signature based on *GBP1*, *IFITM3*, *P2RY14*, and *ID3* was proposed for diagnosing human TB [72]. In our study, *LOC512486*—one of two predicted bovine orthologs of human *GBP1*—was significantly increased in expression after PPD-b stimulation, irrespective of infection status (Supplementary Table 7). In addition, increased expression of *GBP1* was also observed in a gene expression microarray study of PBMC from *M. bovis*-infected cattle stimulated with PPD□b for 12 h [77].

Among the *M. bovis*-infected animals, a core transcriptional response of 170 DE genes was observed (**Fig. 5** and Supplementary Table 8). As part of this signature, in response to *in vitro* stimulation with PPD-b, three interferon-induced GTPase genes (*LOC783920*—predicted to encode an interferon-induced very large GTPase 1-like protein, *GBP4*, and *GBP5*) were increased in expression. These guanylate binding proteins are involved in growth control of intracellular pathogens [78]. The *GBP5* gene also exhibited increased expression in human TB patients versus healthy controls and was included in a broader 15-gene model diagnostic classifier set [72]. Increased expression of seven chemokine genes (*CXCL2*, *CXCL3*, *CXCL5*, *CXCL10*, *XCL1*, *CCL3*, *CCL4*) was also observed in the core transcriptional response for *M. bovis*-infected samples (**Fig. 5** and Supplementary Table 8). Chemokines have key roles in innate immunity and inflammation during mycobacterial infections that lead to tuberculosis disease [79, 80]. Recent work has shown that levels of plasma CCL3 and CXCL10 proteins were significantly elevated in children with active TB [81]. Also, *CXCL10* gene expression and levels of plasma CXCL10 protein has been indicated as a promising biomarker for *M. bovis* infection and bTB disease [82].

**Fig. 5:**
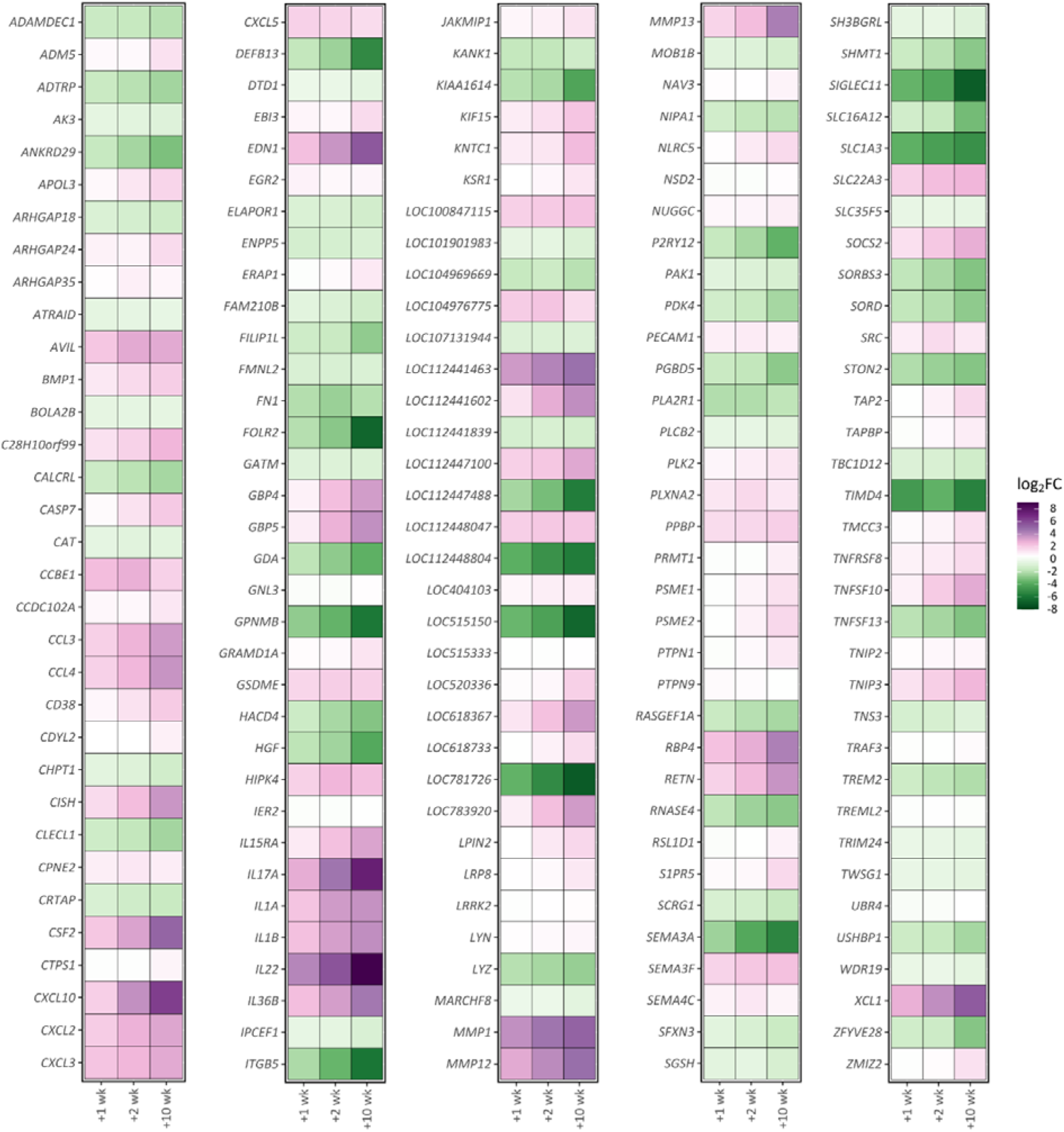
Heatmaps of 170 DE genes across all post-infection time points in response to PPD-b stimulation. Differential gene expression between stimulated and non-stimulated samples is shown for three post-infection time points (+1 wk, +2 wk, and +10 wk) with log_2_ fold-change shown with green to purple heat map colours.

Production of CXCL10 has been found to correlate closely with IFN-γ production [83]. However, CXCL10 alone is not sensitive enough to replace IFN-γ and, as a result, it has been proposed that CXCL10 be used alongside IFN-γ as a dual biomarker [84, 85]. As the gene encoding the cytokine responsible for the hallmark response of bTB surveillance testing, the interferon gamma gene (*IFNG*) was substantially increased in expression in the +2 wk (log_2_FC = + 3.19, FDR *P*_adj._ = 4.54 × 10^-9^) and +10 wk (log_2_FC = + 7.85, FDR *P*_adj._ = 5.05 × 10^-31^) post-infection PPDb-stimulated groups relative to the non-stimulated samples at those time points (Supplementary Tables 5 and 6). In this regard, it is notable that *IFNG* was ranked second by fold-change and third by statistical significance for the +10 wk contrast between the PPDb-stimulated and non-stimulated groups (see **Table 4** and Supplementary Table 6). Taken together, these observations are in concordance with previous work showing that persistent levels of the IFNG cytokine are detectable starting at 2–3 weeks after *M. bovis* infection [15].

The *CXCL8* and *RGS16* genes were upregulated at the +2 wk and +10 wk but not the +1 wk post-infection time points. It is noteworthy, therefore, that these genes have been previously detected in a 19-gene signature of *M. bovis* infection across five post-infection time points in unstimulated peripheral blood samples from the same animals in the experimental time course described here [29]. The *CXCL8* gene (previously known as *IL8*) encodes the interleukin 8 chemokine, a neutrophil chemoattractant that has been shown to be highly expressed in multiple cell types during *M. tuberculosis* infection in vitro and in vivo [79]. Furthermore, polymorphisms in the *CXCL8* gene have been associated with human TB susceptibility [86]. *CXCL8* has also shown to be significantly upregulated after in vitro PPD-b stimulation of *M. bovis*-infected PMBC [87] and monocyte-derived macrophages (MDM) [88]. The *CXCL8* gene was also upregulated in PBL [27] but downregulated in PBMC [89] from *M. bovis*-infected cattle without antigen stimulation. Moreover, the protein encoded by *RGS16* belongs to the regulator of G protein signalling family, members of which have been shown to be dysregulated in multiple diseases [90]. This gene has been implicated in T cell activation, with overexpression in transgenic mice leading to selective blockade of T cell migration to lung parenchyma and induced cellular cytokine synthesis in response to inflammatory stimuli [91]. Upregulation of *RGS16* was also detected in human T cells upon stimulation with IL-2 [92]. Finally, substantially increased expression of both *CXCL8* and *RGS16* has also been observed in bovine alveolar macrophages after in vitro infection with *M. bovis* [93, 94]. In summary, the gene transcriptional signature PPD-b response identified exclusively in the post-infection time points—with particular attention to *CXCL8*, *RGS16*, and *GBP5*—represents a panel of potential biomarkers of *M. bovis* infection and requires further investigation for its use in ancillary diagnostics for bTB surveillance and management in cattle populations.

## 5. Conclusion

High-resolution RNA-seq transcriptional profiling of PPD-b-stimulated peripheral blood collected from *M. bovis*-infected cattle has revealed increasing perturbation of the blood transcriptome across an infection time course up to 10 weeks post-infection. Leukocyte transendothelial migration was observed to be most significantly affected host pathway and core peripheral blood transcriptional responses to PPD-b stimulation were detected that consisted of 93 genes in both non-infected and infected animals across the time course, and 170 genes in infected animals only. This study provides a panel of potential biomarkers for *M. bovis* infection in an accessible tissue amenable to development of novel diagnostics.

## Data availability

The data set presented in this study can be found in online repositories. The names of the repository/repositories and accession number(s) can be found at the European Nucleotide Archive (ENA – www.ebi.ac.uk/ena) with study accession number **PRJEB44568**.

## Author contributions

CNC, BVR, HMV, EG, SVG, and DEM conceived and designed the study and organised bovine sample collection. CNC, JAB, KEM, NCN, DAM, and AOW performed laboratory work including blood sampling, RNA extraction and RNA-seq library generation. CNC, GPM, and DEM performed the statistical analyses. CNC, GPM, and DEM wrote the manuscript. All authors reviewed and approved the final manuscript.

## Acknowledgements

The authors would like to thank all members of the Animal Services Unit of the APHA, Weybridge for their exemplary care of the animals used in these experiments. CNC would like to thank Sarah Faherty O’Donnell for useful discussions and Dr Kerri Malone for assistance with R scripting.

## Supplementary information

Supplementary information for this article can be found online.

