## Supplementary Figures for "High-resolution transcriptomics of bovine purified protein derivative-stimulated peripheral blood from cattle infected with *Mycobacterium bovis* across an experimental time course"

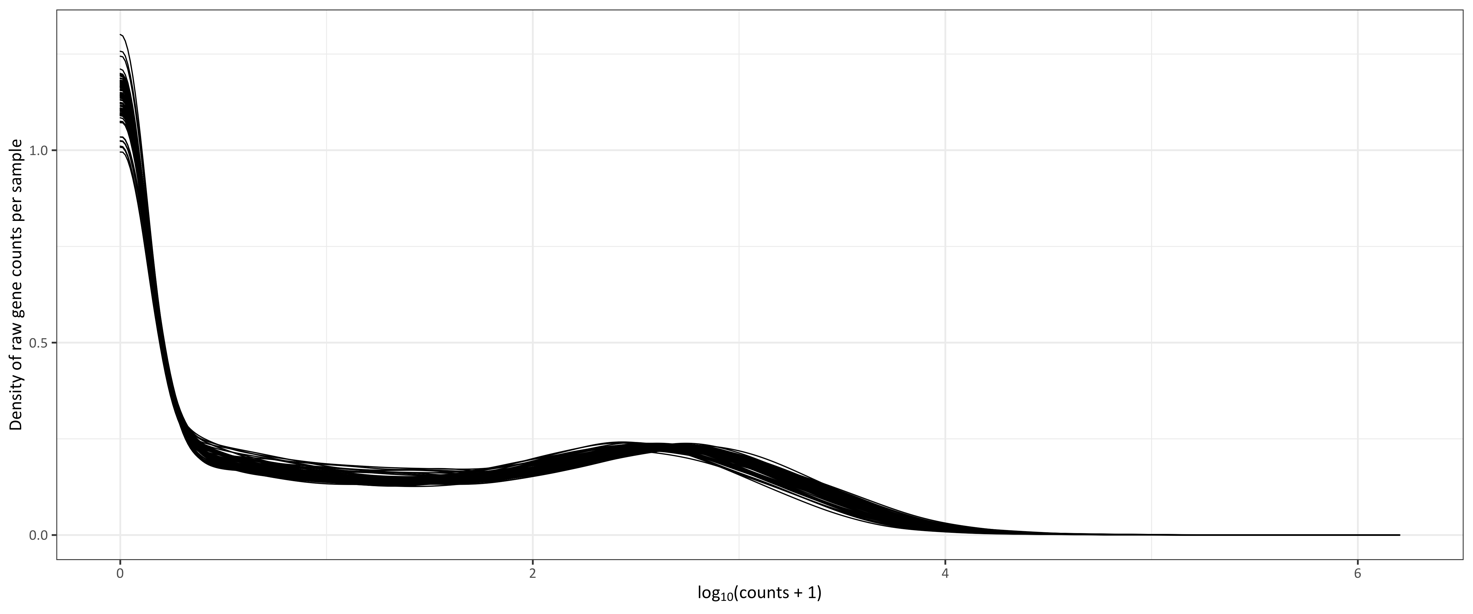

**Supplementary Figure 1:** Density of log_10_ gene counts per library prior to filtering of lowly expressed genes.

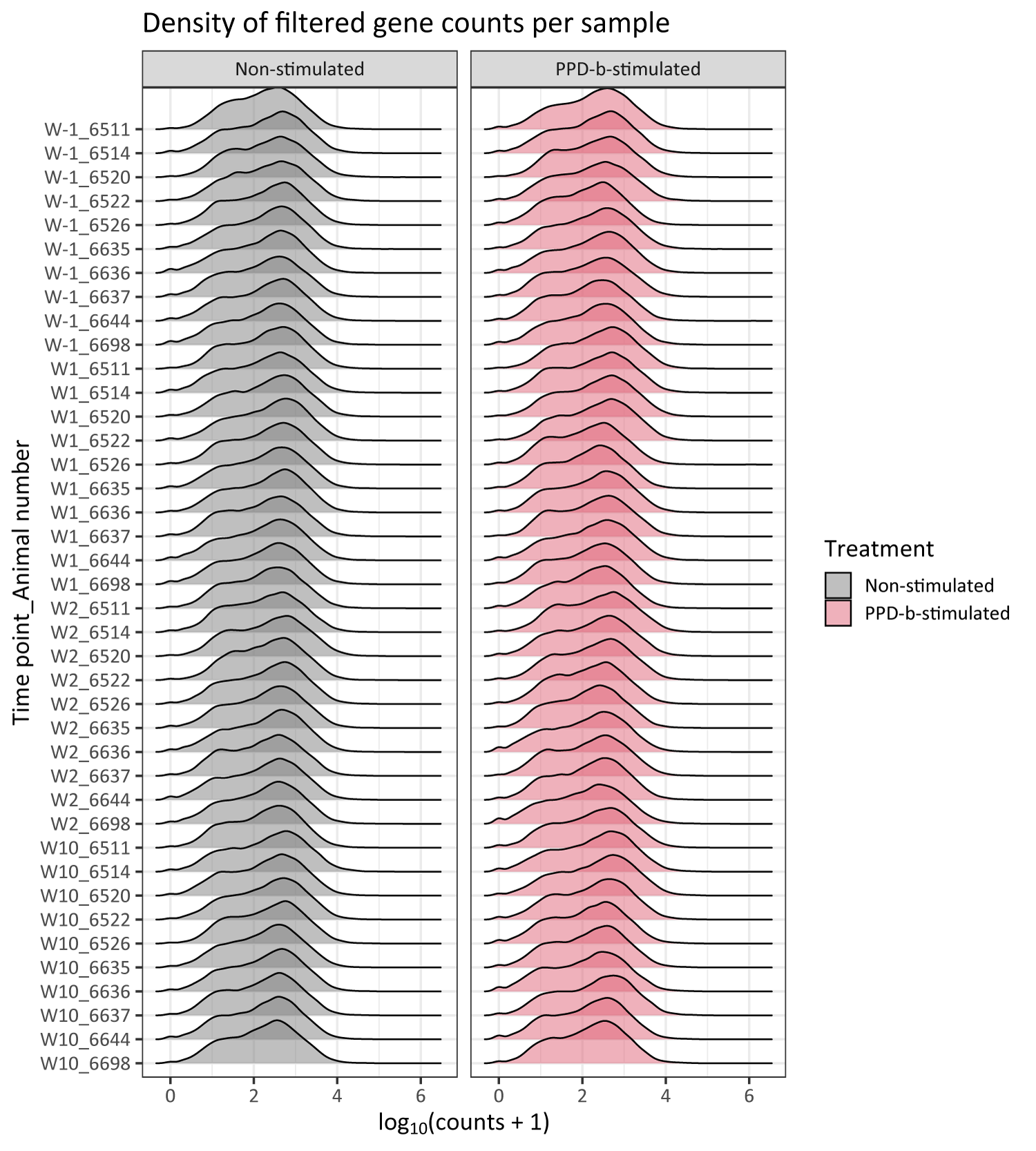

**Supplementary Figure 2:** Ridge plots showing density of log_10_ gene counts per library after filtering of lowly expressed genes.

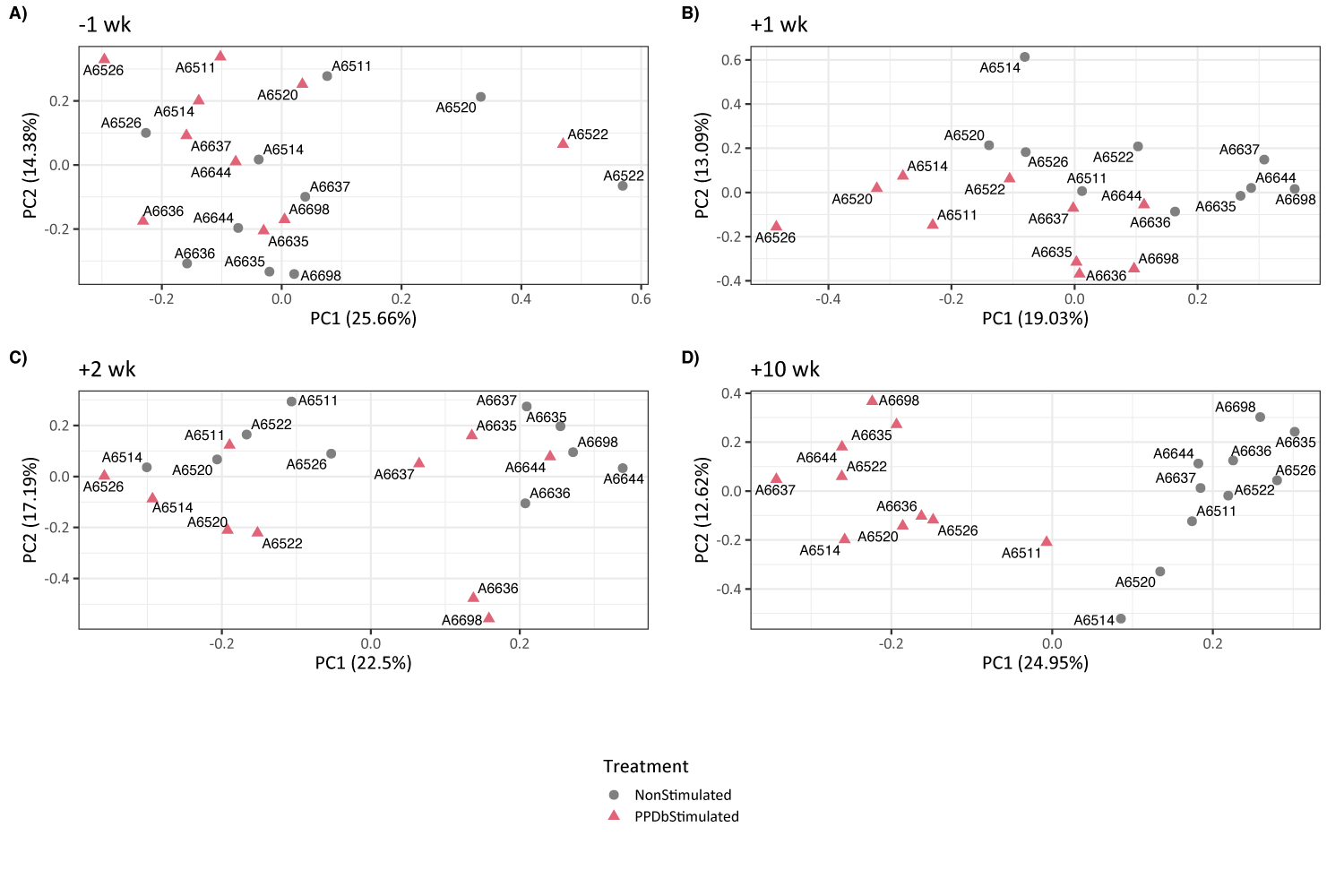

**Supplementary Figure 3:** Principal Components Analysis (PCA) plots generated from RNA-seq expression data (*n* = 20 for each plot).

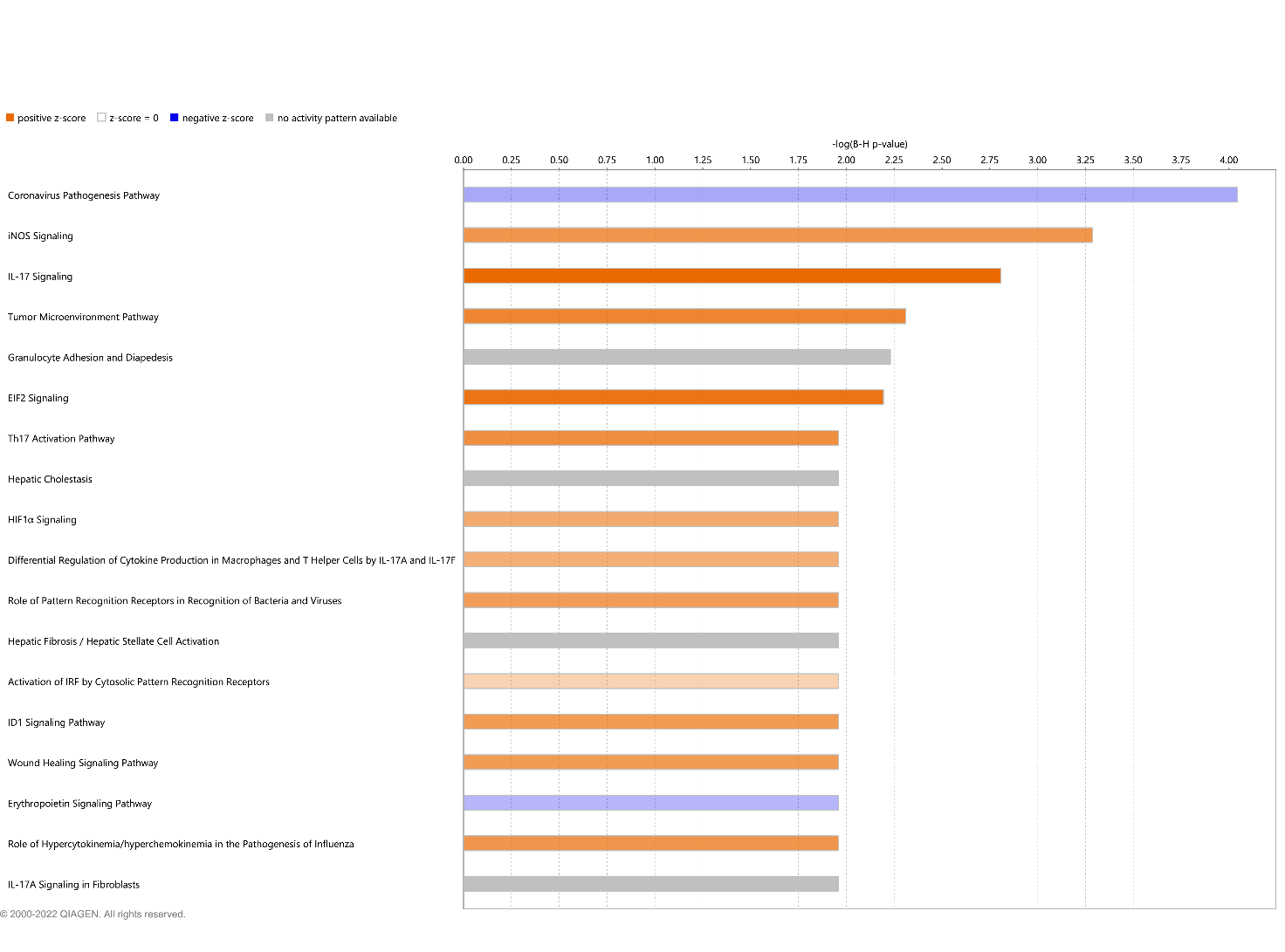

**Supplementary Figure 4:** Bar chart showing the top 18 canonical pathways for the -1 wk time point using IPA^®^ Core Analysis in order of increasing -log(B-H FDR-adjusted *P*-value). The orange and blue colours of the bars indicate the predicted pathway activation or inhibition respectively. Gray bars indicate pathways for which no prediction can be made. In this bar chart the top 18 pathways are shown because the *P*-values are the same as the top 10 pathways to three decimal places.

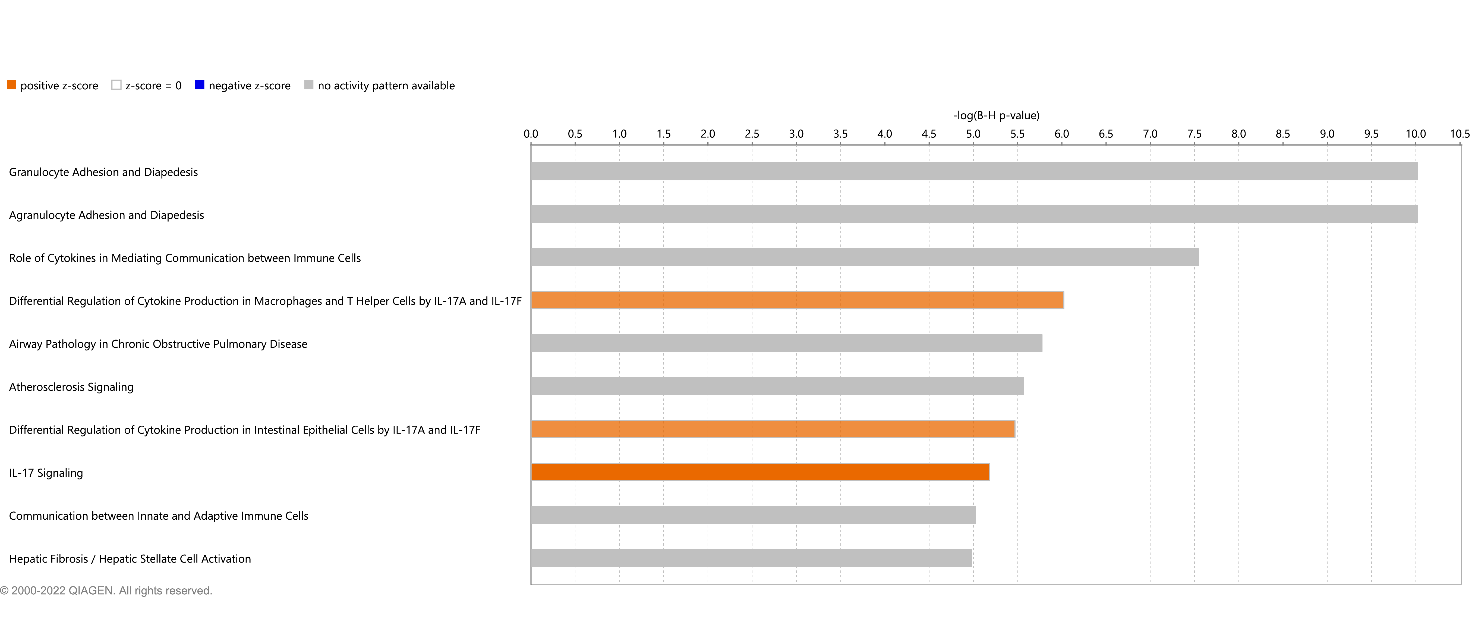

**Supplementary Figure 5:** Bar chart showing the top 10 canonical pathways for the +1 wk time point using IPA^®^ Core Analysis in order of increasing -log(B-H FDR-adjusted *P*-value). The orange and blue colours of the bars indicate the predicted pathway activation or inhibition respectively. Gray bars indicate pathways for which no prediction can be made.

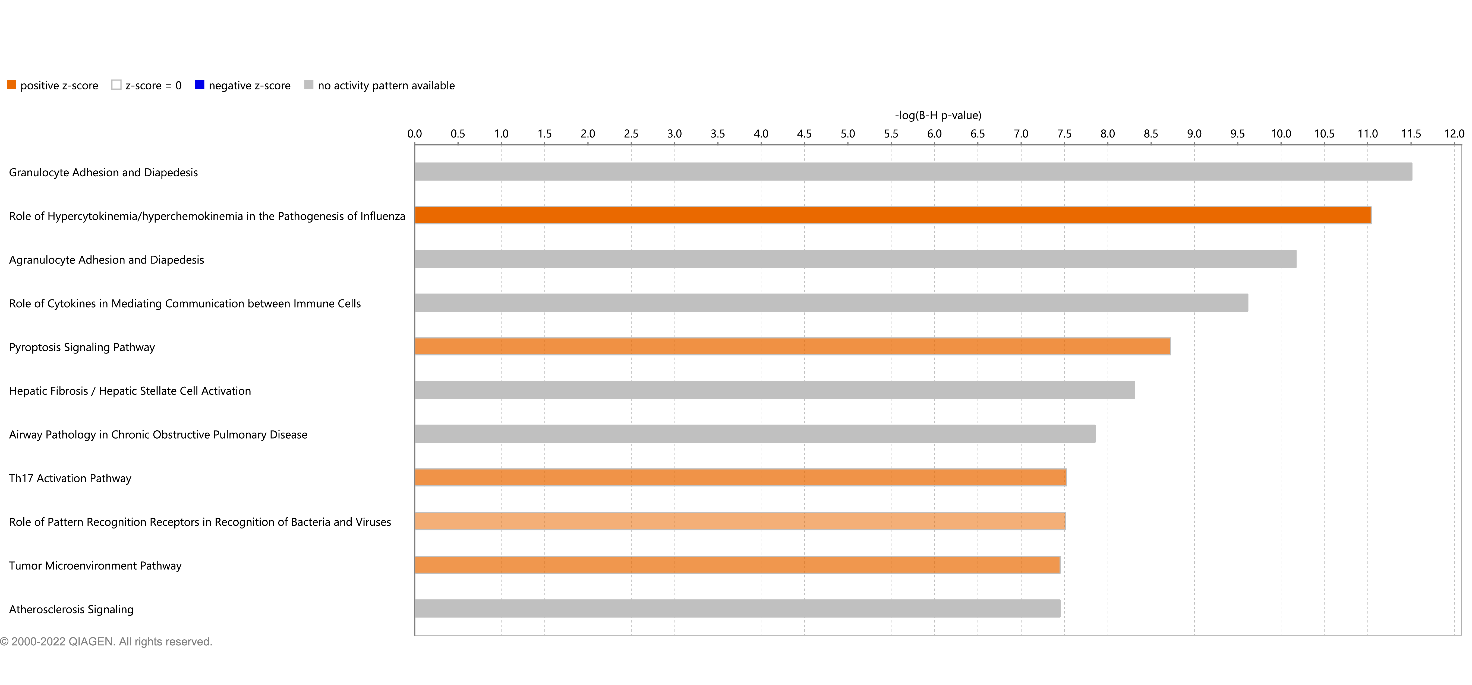

**Supplementary Figure 6:** Bar chart showing the top 11 canonical pathways for the +2 wk time point using IPA^®^ Core Analysis in order of increasing -log(B-H FDR-adjusted *P*-value). The orange and blue colours of the bars indicate the predicted pathway activation or inhibition respectively. Gray bars indicate pathways for which no prediction can be made. In this bar chart the top 11 pathways are shown because the *P*-values are the same as the top 10 pathways to three decimal places.

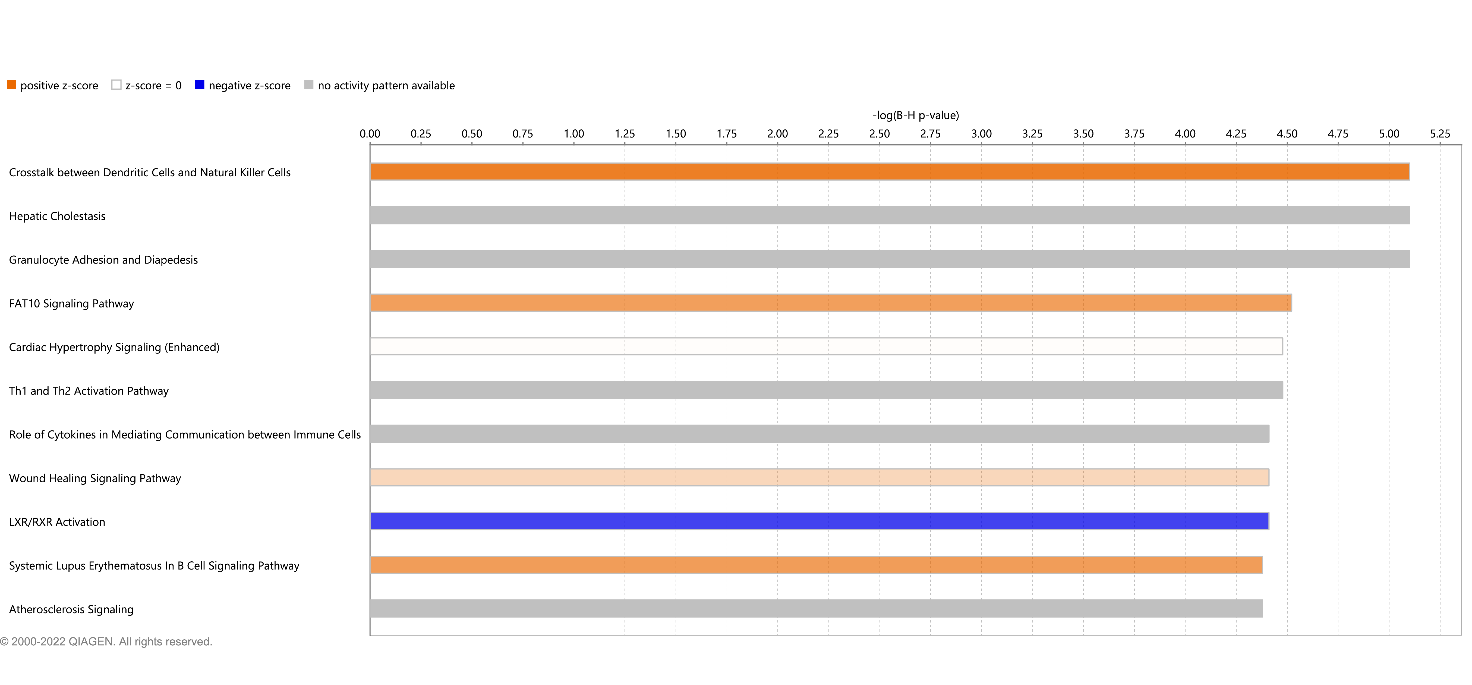

**Supplementary Figure 7:** Bar chart showing the top 11 canonical pathways for the +10 wk time point using IPA^®^ Core Analysis in order of increasing -log(B-H FDR-adjusted *P*-value). The orange and blue colours of the bars indicate the predicted pathway activation or inhibition respectively. Gray bars indicate pathways for which no prediction can be made. In this bar chart the top 11 pathways are shown because the *P*-values are the same as the top 10 pathways to three decimal places.

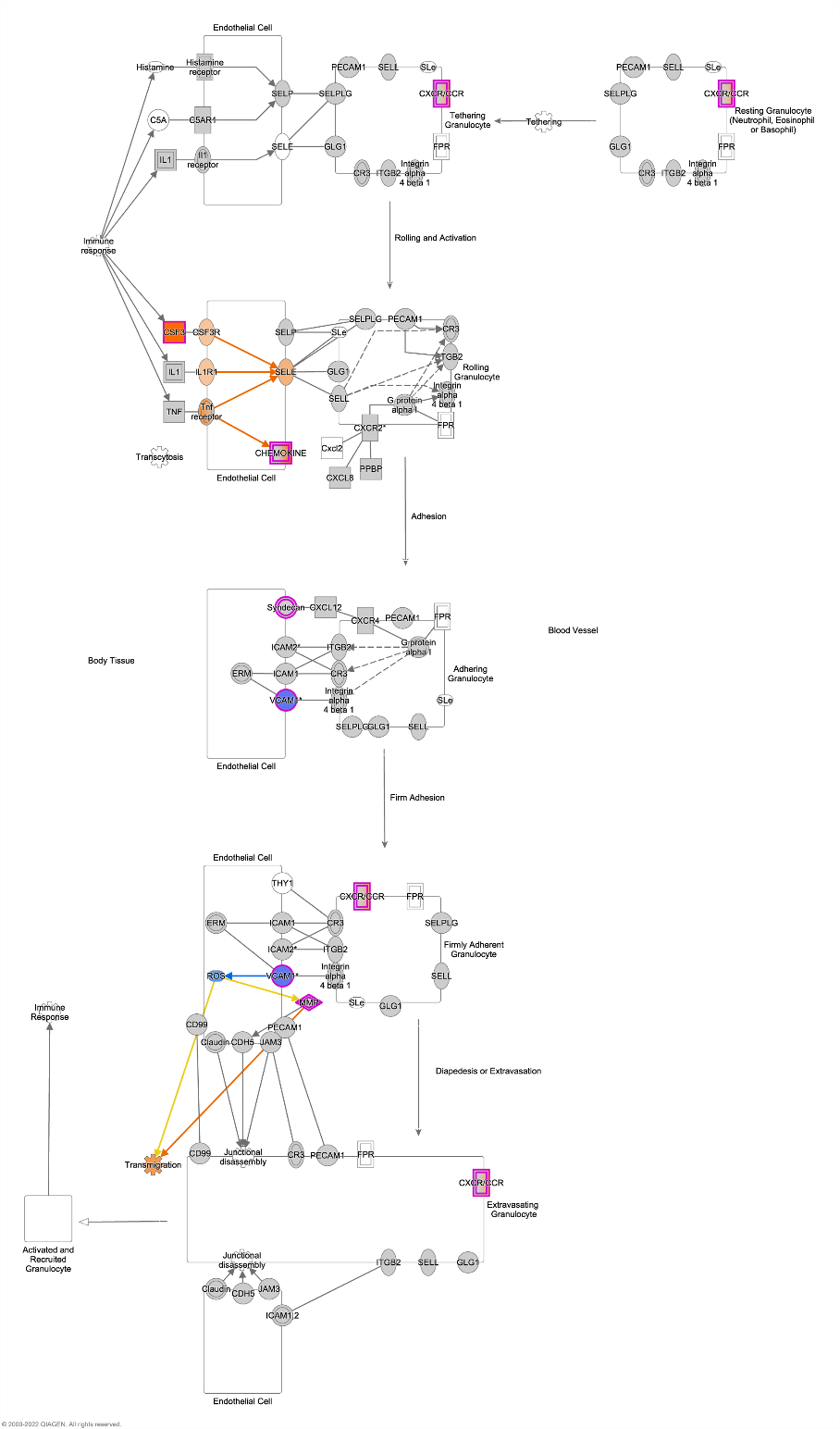

**Supplementary Figure 8: IPA® pathway diagram showing the *Granulocyte Adhesion and Diapedesis* pathway for the -1 wk time point.** Differentially expressed genes in the pathway are highlighted with pink outlines. Increased or decreased expression are indicated by shades of orange and blue respectively. Genes with a double outline containing a gradient of these colours indicate groups or complexes of genes that may be significantly perturbed. A legend is available at <https://qiagen.secure.force.com/KnowledgeBase/articles/Basic_Technical_Q_A/Legend>.

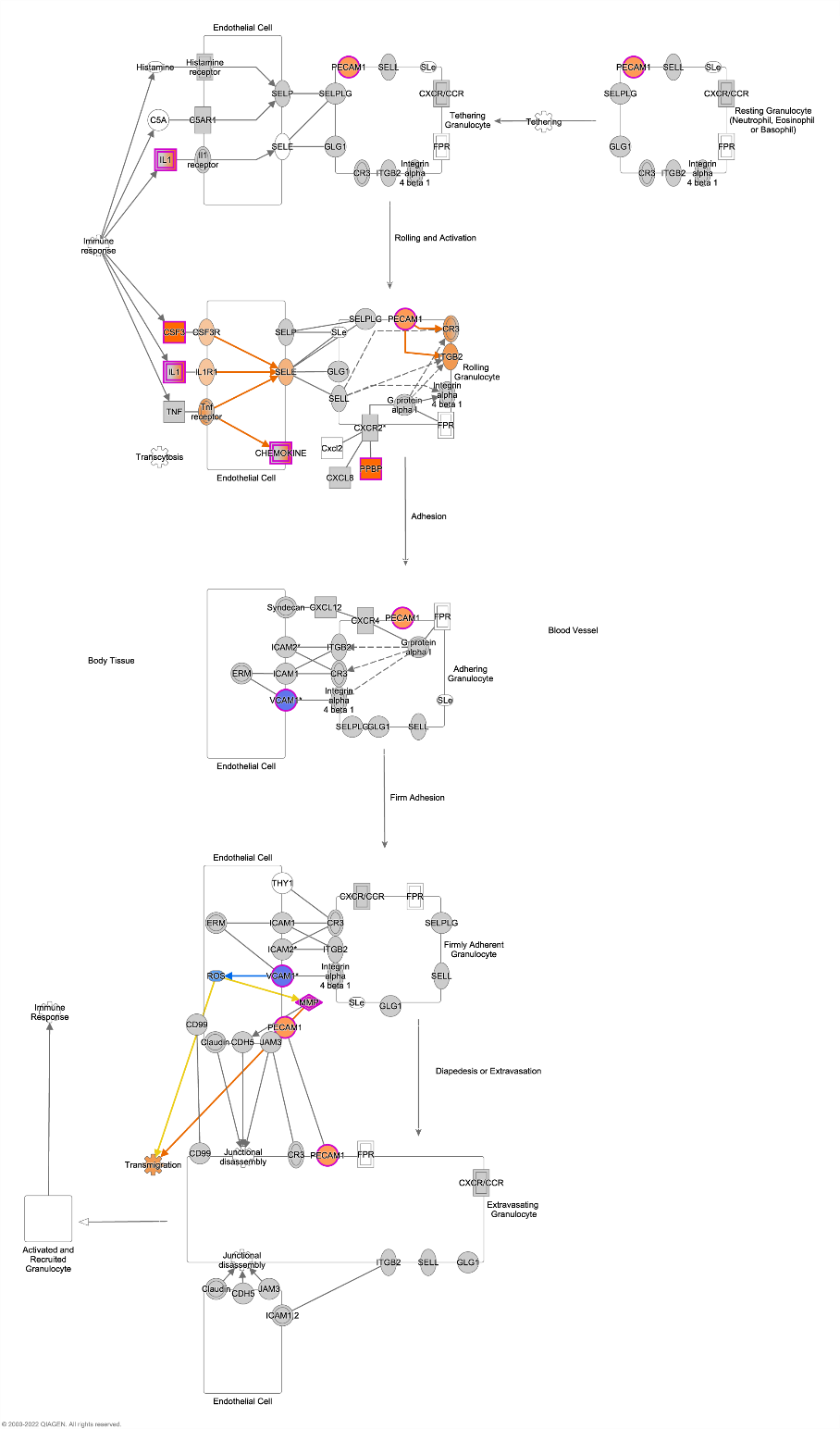

**Supplementary Figure 9: IPA® pathway diagram showing the *Granulocyte Adhesion and Diapedesis* pathway for the +1 wk time point.** Differentially expressed genes in the pathway are highlighted with pink outlines. Increased or decreased expression are indicated by shades of orange and blue respectively. Genes with a double outline containing a gradient of these colours indicate groups or complexes of genes that may be significantly perturbed. A legend is available at <https://qiagen.secure.force.com/KnowledgeBase/articles/Basic_Technical_Q_A/Legend>.

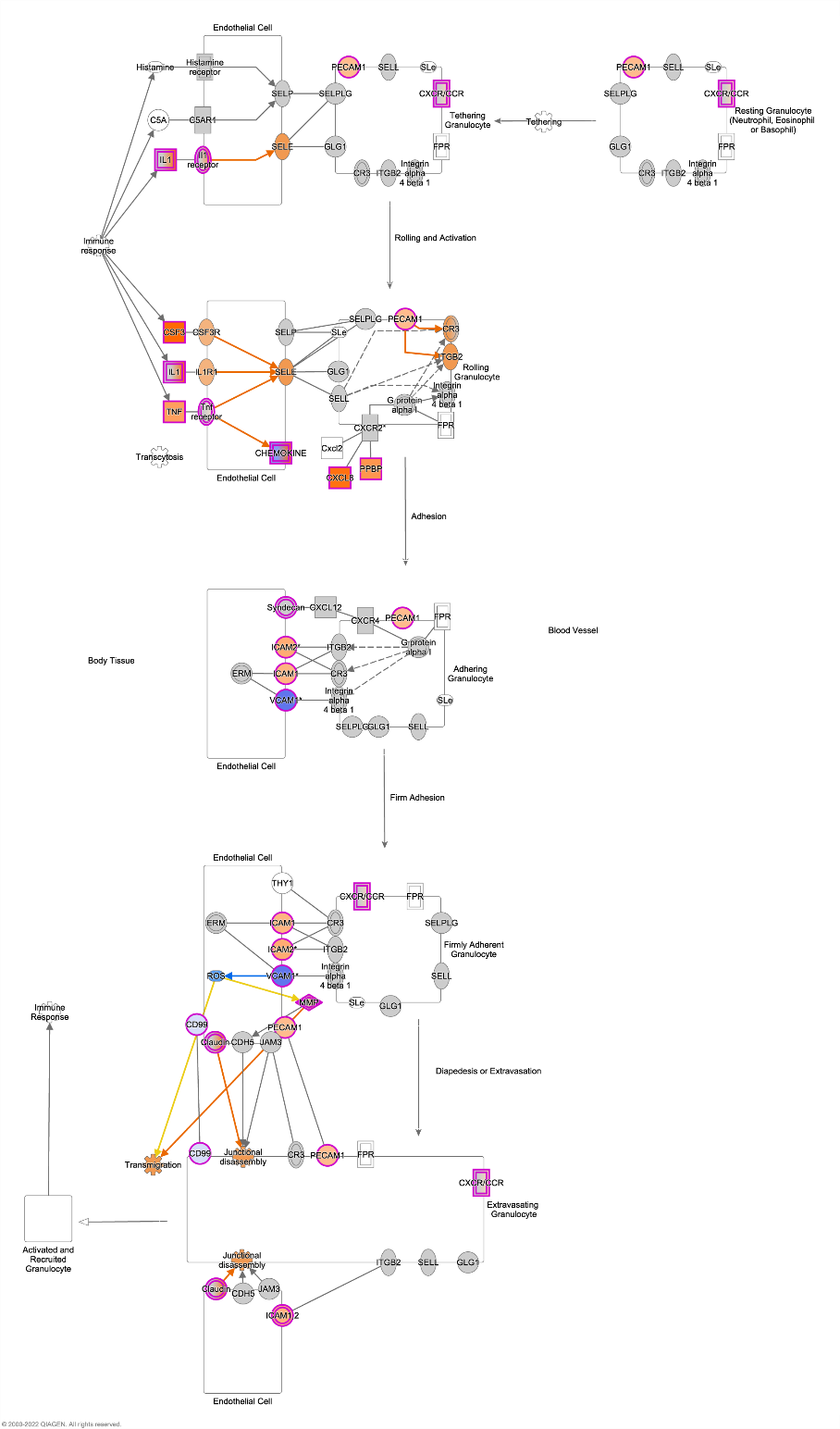

**Supplementary Figure 10: IPA® pathway diagram showing the *Granulocyte Adhesion and Diapedesis* pathway for the +2 wk time point.** Differentially expressed genes in the pathway are highlighted with pink outlines. Increased or decreased expression are indicated by shades of orange and blue respectively. Genes with a double outline containing a gradient of these colours indicate groups or complexes of genes that may be significantly perturbed. A legend is available at <https://qiagen.secure.force.com/KnowledgeBase/articles/Basic_Technical_Q_A/Legend>.

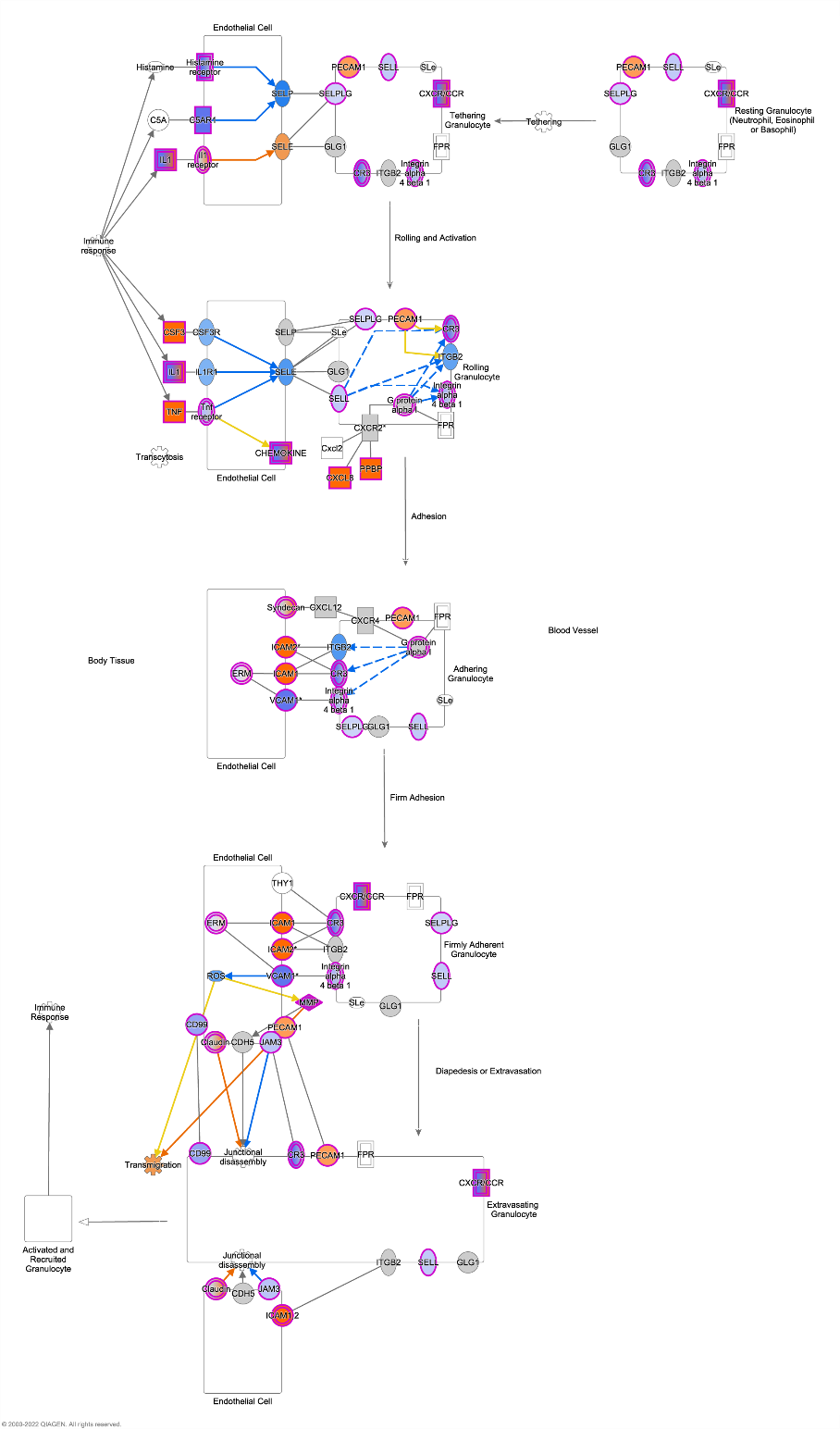

**Supplementary Figure 11: IPA® pathway diagram showing the *Granulocyte Adhesion and Diapedesis* pathway for the +10 wk time point.** Differentially expressed genes in the pathway are highlighted with pink outlines. Increased or decreased expression are indicated by shades of orange and blue respectively. Genes with a double outline containing a gradient of these colours indicate groups or complexes of genes that may be significantly perturbed. A legend is available at <https://qiagen.secure.force.com/KnowledgeBase/articles/Basic_Technical_Q_A/Legend>.
